# Storm: Incorporating transient stochastic dynamics to infer the RNA velocity with metabolic labeling information

**DOI:** 10.1101/2023.06.21.545990

**Authors:** Qiangwei Peng, Xiaojie Qiu, Tiejun Li

## Abstract

The time-resolved scRNA-seq (tscRNA-seq) provides the possibility to infer physically meaningful kinetic parameters, e.g., the transcription, splicing or RNA degradation rate constants with correct magnitudes, and RNA velocities by incorporating temporal information. Previous approaches utilizing the deterministic dynamics and steady-state assumption on gene expression states are insufficient to achieve favorable results for the data involving transient process. We present a dynamical approach, Storm (Stochastic models of RNA metabolic-labeling), to overcome these limitations by solving stochastic differential equations of gene expression dynamics. The derivation reveals that the new mRNA sequencing data obeys different types of cell-specific Poisson distributions when jointly considering both biological and cell-specific technical noise. Storm deals with measured counts data directly and extends the RNA velocity methodology based on metabolic labeling scRNA-seq data to transient stochastic systems. Furthermore, we relax the constant parameter assumption over genes/cells to obtain gene-cell-specific transcription/splicing rates and gene-specific degradation rates, thus revealing time-dependent and cell-state specific transcriptional regulations. Storm will facilitate the study of the statistical properties of tscRNA-seq data, eventually advancing our understanding of the dynamic transcription regulation during development and disease.

## Background

Cells are dynamic identities that are subject to intricate transcriptional and post-transcriptional regulations. Understanding the tight regulation of the RNA life cycle will shed light on not only the regulatory mechanism of RNA biogenesis, but also cell fate transitions. Based on the observation that most scRNA-seq approaches capture both premature unspliced mRNA and mature spliced mRNA information, La Manno et al. La Manno et al. (2018) pioneered the concept of RNA velocity or the time derivative of spliced RNA to reveal the local fate of each individual and designed a RNA kinetic parameter inference method called velocyto based on the steady state assumption. In a later work, scVelo Bergen et al. (2020) relaxed the steady-state assumption and proposed a dynamic RNA velocity model to infer gene-specific reaction rates of transcription, splicing and degradation as well as cell-specific hidden time using the expectation-maximization (EM) algorithm. Li et al. Li et al. (2021) derived a stochastic model of RNA velocity based on the chemical master equation (CME) satisfied by the probabilistic mass function (PMF) rather than the deterministic ordinary differential equation (ODE) satisfied by the mean, and presented a mathematical analysis framework of RNA velocity. MultiVelo Li et al. (2022) extends the dynamic RNA velocity model by incorporating epigenome data that can be jointly measured with emerging multi-omics approaches. Protaccel Gorin et al. (2020) extends the concept of RNA velocity to protein. UniTVelo Gao et al. (2022) uses a top-down design for more flexible estimation of the RNA velocity, as opposed to the usual bottom-up strategy. DeepVelo Cui et al. (2022) uses graph convolutional neural networks to infer cell-specific parameters to extend RNA velocity to cell populations containing time-dependent dynamics and multiple lineages which were proven to be challenging in previous methods Bergen et al. (2021). Other deep learning-based approaches include VeloVI Gayoso et al. (2022), VeloVAE Gu et al. (2022), LatentVelo Farrell et al. (2022), cellDancer Li et al. (2023), and so on. However, due to the absence of physical time information, the above methods usually suffer the issue of scale invariance, that is, amplifying the parameters by an arbitrary amount will not change the solution if the time shrinks with the same amount, e.g., the exact physical time remains undetermined. This issue makes the inferred parameters and the RNA velocity have physical significance only up to a multiplicative constant Li et al. (2021). In addition, the missing time information enters the model as hidden variables, which makes the parameter inference difficult.

Technological innovations in scRNA-seq now enable us to directly measure the amount of newly synthesized mRNA molecules over a short period of time, either through chemically introduced mutations in the sequencing reads or direct biotin pull-down of RNA analogs such as 4sU metabolically labeled RNA molecules, which subtly introduces physical time information. These time-resolved metabolic labelingaugmented scRNA-seq (tscRNA-seq) include scSLAM-seq Erhard et al. (2019), scNT-seq Qiu et al. (2020), sci-fate Cao et al. (2020), NASC-seq Hendriks et al. (2019) and scEU-seq Battich et al. (2020). Qiu et al. Qiu et al. (2022) recently developed Dynamo to reconstruct analytical vector fields from discrete RNA velocity vectors by taking advantage of tscRNA-seq data to infer more robust and time-resolved RNA velocity, however, they only used the deterministic model and largely relied on the steady-state assumption.

To overcome the shortcomings of Dynamo and fully explore the potential of tscRNA-seq data, we present the Storm approach (Stochastic models of RNA metabolic-labeling) to improve the estimation of RNA kinetic parameters and the inference of the RNA velocity of the metabolic labeling scRNA-seq data by incorporating the transient stochastic dynamics of gene expressions. Importantly, we focus on modeling the kinetics/pulse metabolic labeling data as it follows the RNA synthesis across multiple short time periods and is thus ideal to capture temporal RNA kinetics. In order to properly model both biological noise and cell-specific technical noise (due to the variations in sequencing depth across individual cells and dropout resulting from imperfect RNA capture in scRNA-seq), we implemented in Storm three stochastic models of new mRNA (or new unspliced and spliced mRNA). Depending on the biological processes considered, Storm indicates that new mRNA sequencing data obeys different types of cell-specific Poisson (CSP) distributions. On this basis, Storm also includes hypothesis testing, parameter inference and goodness of fit evaluation methods for CSP-type distribution. In addition, we analyze the similarities and differences of the model considering RNA splicing or not. For one-shot data, we introduce the steady-state assumption to make the parameter inference possible. We verified the effectiveness of Storm in the cell cycle data set of kinetic experiments from the scEU-seq study Battich et al. (2020) and several one-shot datasets, including scSLAM-seq, scNT-seq and scifate. Storm is incorporated in Dynamo Qiu et al. (2022) of the Aristotle ecosystem that facilitates rich downstream analytical vector field modeling.

## Results

### Overall description of Storm

We established three stochastic gene expression models for new mRNA (or new unspliced and spliced mRNA) (**Fig. 1A**) for the inference of the RNA kinetic parameters and thus the RNA velocity in the Storm approach. In Model 1, only transcription and mRNA degradation were considered. In Model 2, we considered transcription, splicing, and spliced mRNA degradation. And in Model 3, we considered the switching of gene expression states, transcription in the active state, and mRNA degradation.

**Figure 1:**
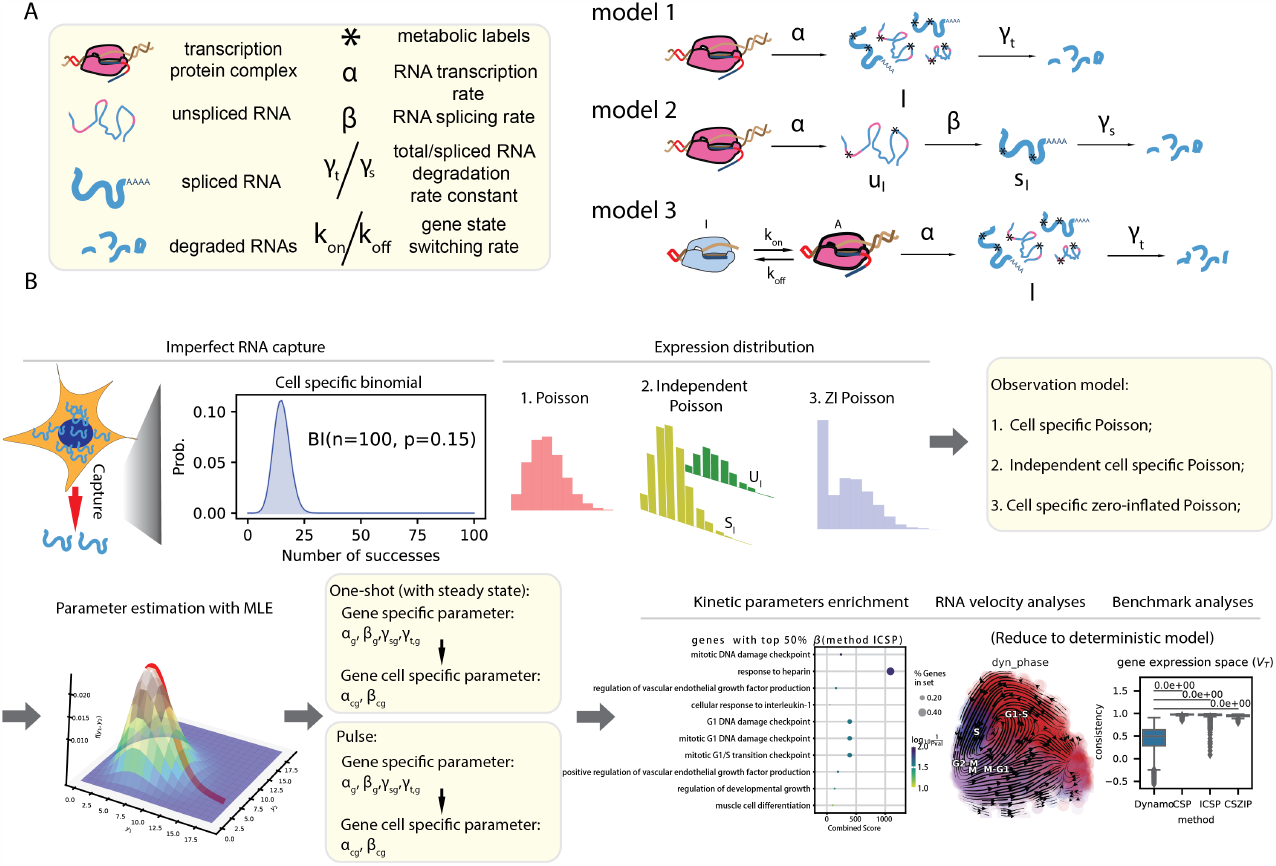
Schematic overview of Storm. **A**. Three models of RNA life cycle considering different biological processes: **Model 1**: Reaction dynamics model for new RNA *l*(*t*) ignoring the splicing process, where *α* is the transcription rate and *γ*_*t*_ is the total mRNA degradation rate. **Model 2**: Reaction dynamics model of new unspliced and spliced mRNA (*u*_*l*_(*t*), *s*_*l*_(*t*)) considering the splicing process, where *β* is the splicing rate, *γ*_*s*_ is the spliced mRNA degradation rate, and *α* is the same as **Model 1**. Reaction dynamics model of new RNA *l*(*t*) considering gene state switching, where *α* and *γ*_*t*_ are the same as in **Model 1**, *k*_on_ is the rate at which the gene switches from the inactive state to the active state, *k*_off_ is the opposite. **B**. Complete workflow diagram for parameter inference and downstream analysis based on stochastic dynamics of new mRNA considering technical noise.

The complete workflow of Storm is demonstrated in **Fig. 1B**. We first analytically solve the new RNA (or new unspliced and spliced mRNA) stochastic dynamics corresponding to the above three models, which are Poisson distribution, independent Poisson distribution and zero-inflated Poisson distribution, respectively. In addition, we model the technical noise as the cell-specific binomial distribution. By integrating the biological noise and the technical noise together, we obtain the distribution for the measured number of new/labeled mRNA molecules (or new unspliced and spliced mRNA molecules), which are cell-specific Poisson distribution, independent cell-specific Poisson distribution and cell-specific zero-inflated Poisson distribution, respectively. Maximum likelihood estimation (MLE) is used to fit the data and make inferences for the parameters shown in the corresponding models.

To ensure the general applicability of Storm in common nascent RNA labeling schemes, such as one-shot or kinetics/pulse experiments (See Figure 2 of Qiu, et. al Qiu et al. (2022)), we designed specific estimation strategies for each labeling scheme. For the one-shot labeling experiments, since there is only one labeling duration, the steady-state assumption under the stochastic dynamics framework is reinvoked to infer parameters. For kinetics/pulse-labeling experiments with multiple labeling durations, the transient stochastic dynamics framework is used without the steady-state assumption. Furthermore, the goodness-of-fit index 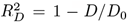 based on deviance commonly used in generalized linear models is utilized to quantify the goodness of fit of our models in kinetics/pulse datasets. The index is then used to select genes that are more consistent with model assumptions for later downstream analysis, such as the enrichment analysis of different gene-specific parameters. Furthermore, we relaxed the previous assumption of constant parameters in genes or cells and assumed that only degradation rates (*γ*_*t*_ in Models 1 and 3; *γ*_*s*_ in Model 2) are constant while the other parameters (*α* in three models; *β* in Model 2; *p*_off_ in Model 3) are cell specific and depend on the state of gene expression in each cell. This relaxation would be useful for modeling lineage-specific kinetics resulted from hierarchical lineage bifurcation, which is common in cell developments. Finally, in order to calculate and visualize the RNA velocity, we reduced the considered stochastic models to derive the deterministic equation for the mean gene expression. The inferred parameters, after filtering with the goodness-of-fit index are then used in RNA velocity analysis and visualization. Notably, to demonstrate Storm’s performance, we conducted systematic comparison with the state-of-the-art method Dynamo Qiu et al. (2022) for processing metabolic labeling scRNA-seq experiment datasets.

**Figure 2:**
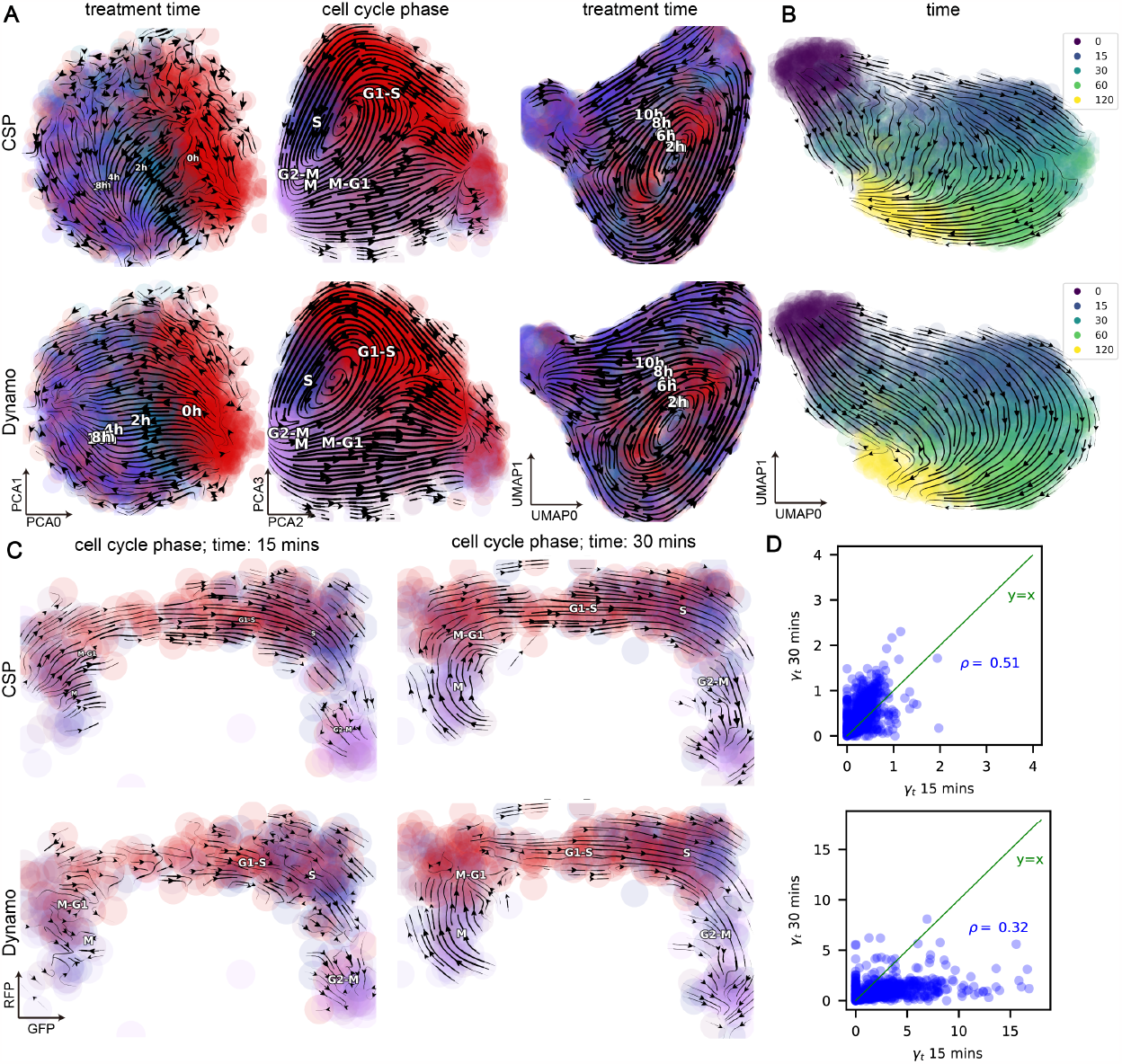
Stochastic model combined with steady-state assumptions for one-shot experiments. **A**. Streamline plots of the sci-fate dataset Cao et al. (2020) reveal two orthogonal processes of GR response and cell-cycle progression. From left to right: streamline plot on the first two PCs, the second two PCs, and the first two UMAP components that are reduced from the four PCs, respectively. The first row is the result of CSP and the second row is the result of Dynamo. The same applies to panels **B, C**, and **D. B**. Streamline projected in the UMAP space plots of neuronal activity under KCl polarization datasets from scNT seq Qiu et al. (2020). **C**. Streamline projected in the RFP_GFP space plots of cell cycle dataset from scEU-seq Battich et al. (2020). On the left is the result of taking only the data labelled with 15 minutes, and on the right is the data labelled with 30 minutes. **D**. Comparison of degradation rates *γ*_*t*_ in cell cycle datasets with labeling duration of 15 and 30 minutes.

In the continued subsections we will present the details of each step in the Storm workflow, starting from the introduction of our mathematical models.

### CSP modeling of counts data with metabolic labeling information

We proposed and analytically solved three aforementioned stochastic gene expression models for the dynamics of new mRNAs (or new unspliced and spliced mRNAs).

For simplicity of modeling, we followed La Manno et al. (2018); Bergen et al. (2020) to assume that the genes are independent. In the stochastic gene expression model, the generation of new/labeled mRNA 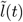 (or new unspliced and spliced mRNA 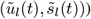 is a stochastic process, and we are interested in the evolution of its PMF over time,which is denoted by

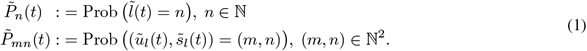

In Model 1 and Model 2, since the initial value of 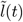 (or 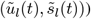) is 0, we obtained the following closed-form solution (see “Methods” section).

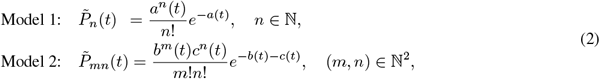

where

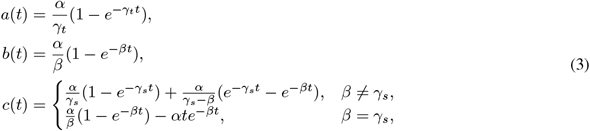

which means that 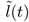 obeys the Poisson distribution with mean *a*(*t*) in Model 1, and 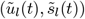 obey independent Poisson distributions with mean *b*(*t*) and *c*(*t*) in Model 2. Here *α, β* are the transcription and splicing rates, and *γ*_*s*_, *γ*_*t*_ are the spliced and total mRNA degradation rates, respectively.

In Model 3, following Chong et al. (2014), we assumed that switching rates *k*_on_ and *k*_off_ are significantly smaller than *α* and *γ*_*t*_, which implies that the gene expression is either always on or always off during transcription/degradation period. Therefore, 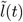 obeys a zero-inflated Poisson (ZIP) distribution, then we have

Model 3:

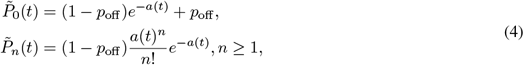

where *p*_off_ = *k*_off_ */*(*k*_on_ + *k*_off_) is the probability that gene expression is in the off state, i.e., the extra proportion of zeros in the ZIP distribution (see “Methods” section).

We also specifically modeled technical noise of the measured number of new RNA (or new unspliced and spliced mRNA) molecules in scRNA-seq experiments. Such noises often lead to dropout of RNA measurements during the sequencing process and generally result in variations in sequencing depth across cells. To account the noise, in Storm we modeled the dropout process of sequencing technology as cell-specific binomial distributions. Adopting the common practice in many preprocessing pipelines through a size factor to normalize the data La Manno et al. (2018); Bergen et al. (2020); Cui et al. (2022); Gayoso et al. (2022); Qiu et al. (2022), we assumed that the total numbers of mRNA molecules across all genes in different cells are close. Probabilistically, this assumption implies that

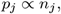

where *p*_*j*_ is the probability of mRNA molecules being captured in cell *j* and *n*_*j*_ is the total number of mRNA molecules across all genes in cell *j* in scRNA-seq experiments.

Combining the stochastic models for biological and technical noise, we can obtain different formalism of the distribution for the measured number of new/labeled mRNA molecules *l*(*t*) (or new unspliced and spliced mRNA molecules (*u*_*l*_(*t*), *s*_*l*_(*t*))) in the scRNA-seq experiments (see “Methods” section) for each model. Specifically, in Model 1, *l*(*t*) obeys the cell-specific Poisson (CSP) distribution, that is,

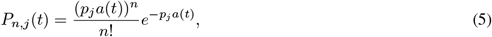

where *P*_*n,j*_ (*t*) is the PMF for the measured number of new mRNA molecules in cell *j*. In Model 2, (*u*_*l*_(*t*), *s*_*l*_(*t*)) obeys the independent cell-specific Poisson (ICSP) distribution, that is,

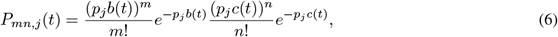

where *P*_*mn,j*_ (*t*) is the joint PMF for the measure number of new unspliced and spliced mRNA molecules in cell *j*. In Model 3, *l*(*t*) obeys the cell-specific zero-inflated Poisson (CSZIP) distribution, that is,

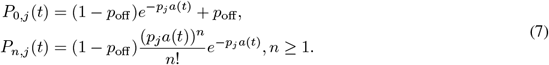

We call the above distributions as *cell-specific* because different cells obey the distributions with different parameters.

Note that Grün et al. also modeled the scRNA-seq data by integrating biological noise and technical noise Grün et al. (2014). Our work is different from them in the following aspects: (1) Our work models the transient dynamics of new mRNA and solves their distribution for the proposed stochastic models analytically. However, in Grün et al. (2014), they instead modeled the total mRNA and derived that the biological noise follows a negative binomial distribution as the steady state of the transcriptional bursting model. (2) Our work accurately models the technical noise as a cell-specific binomial distribution, while they approximated the cell-sepcific binomial distribution with a Poisson distribution and modeled the capture probability as a random variable subject to the Gamma distribution, which finally leads to a negative binomial distribution (Poisson-Gamma mixture distribution) of the technical noise.

As one-shot labeling experiments are much more convenient than pulse experiments in practice, in the following, we will first demonstrate how Storm can be applied to the one-shot case. We will then extensively show Storm’s power in analyzing the pulse datasets.

### Stochastic models combined with steady-state assumptions for one-shot data

Since one-shot data has only one labeling duration, we designed the corresponding parameter inference method which invokes the steady-state assumption under the stochastic model, focusing specifically on Model 1 (see “Methods” section). Similar steady-state methods of the stochastic model can also be designed for both Model 2 and Model 3 as well, although they are not the focus of this paper.

We validated our method in several one-shot datasets (**Fig. 2**, S1). We first analyzed a dataset from the sci-fate study Cao et al. (2020) in which cell cycle progression and glucocorticoid receptor (GR) activation were explored. Similar to Dynamo, the RNA velocity flow from our method also revealed a sequential transition of cells following the DEX (dexamethasone) treatment times in the first two principal components (PCs) (**Fig. 2A Left**). In the second two PCs, we observed an orthogonal circular progression of the cell cycle (**Fig. 2A Middle**). From the first two UMAP dimensions projected further from the four PCs, we observed a combined dynamics of GR responses and cell cycle progression (**Fig. 2A Right**). Next, we analyzed the neuronal activity dataset from the scNT-seq study Qiu et al. (2020) to investigate cellular polarization dynamics after KCl treatment (**Fig. 2B**). Dynamo and Storm both revealed a coherent transition that nicely follows the temporal progression from time point 0 to 15, 30, 60 and finally 120 minutes. We analyzed the murine intestinal organoid system dataset from scEU-seq Battich et al. (2020). Dynamo observed a bifurcation (Fig. S1B, top row) from intestinal stem cells into the secretory lineage (left) and the enterocyte lineage (right), and Storm also observed similar results, although with some defects in the secretory lineage (Fig. S1B, bottom row). We also analyzed mouse fibroblast cells dataset from scSLAM-seq Erhard et al. (2019). We observed that both Dynamo and Storm inferred velocities further discriminated infected from non-infected cells (**Fig. S1C**).

To demonstrate the precision and robustness of the Storm method in estimating the one-shot dataset, we bench-marked the estimated kinetic parameters of different subsets of the cell cycle pulse-labeling dataset Battich et al. (2020), each with a different duration of labeling. On the 15-minute labeling sub-dataset, Storm recovers a transition that matches well with the cell-cycle progression, while the transition recovered by Dynamo is problematic near the M/M-G phase (**Fig. 2C Left**). On the 30-minute labeling sub-dataset, both methods recover the cell cycle progression correctly, but the streamlines of our method are considerably smoother compared to those of Dynamo (**Fig. 2C Right**). In addition, we compared the consistency of degradation rates *γ*_*t*_ inferred by the two methods between two sub-datasets with different labeling durations (**Fig. 2D**). The results show that our method is more consistent compared with Dynamo. Notably, although Storm shows higher consistency than Dynamo, it is still not satisfactory, perhaps due to the experimental noises from different labeling durations and the violation of the steady-state assumption. Therefore, it is crucial to integrate data of different durations of labeling when a kinetic experiment is available. Furthermore, it is equally important to design methods that do not rely on the steady-state assumption for parameter inference.

Finally, we quantitatively compared the degradation rates *γ*_*t*_ inferred by the two methods. The two methods are close on the other datasets (**Fig. S1A**,**D**) except on 15-minute labeling cell cycle sub-dataset where Dynamo is unreasonably large (**Fig. S1A, third column**). Thus, our method has similar or even better performance compared to Dynamo on the one-shot dataset.

### Statistical analysis of cell cycle dataset based on Storm’s stochastic model

Next we first performed a goodness-of-fit test of the stochastic model proposed in Storm to a cell cycle dataset from scEU-seq Battich et al. (2020) with mutliple labeling time points to validate our proposals.

When the fixed labeling duration is *t*_fixed_, *a*(*t*_fixed_), *b*(*t*_fixed_) and *c*(*t*_fixed_) are all fixed constants. We can test whether the number of new mRNA molecules in tscRNA-seq within a fixed labeling duration matches the distribution obtained based on the stochastic models (Eqs. (5), (6) and (7)), respectively. A common method of testing whether a dataset obeys a given distribution is the chi-square (*χ*^2^) goodness-of-fit test Pearson (1900). However, the usual *χ*^2^ test is not directly applicable because in our case different cells obey different distributions with different parameters. By inspecting the mathematical analysis procedure of the *χ*^2^ test Benhamou and Melot (2018), we constructed a new asymptotic *χ*^2^ statistics and proposed a modified *χ*^2^ test for our cell-specific distributions (see “Methods” section).

We used the proposed cell-specific *χ*^2^ test in the cell cycle dataset from the scEU-seq study Battich et al. (2020), in which cells were labeled for 15, 30, 45, 60, 120 or 180 minutes. Because the labeled unspliced mRNA counts *u*_*l*_(*t*) were too small to be grouped/binned to create a distribution, hypothesis tests were performed only for CSP and CSZIP distributions and not for ICSP. The results are shown in **Table 1**. We found that some genes were not well determined (especially for cases with a short duration of labeling) in the sense that these genes had too few new mRNA molecules in the tscRNA-seq experiments, which results in very few groupings and perfect fittings. With so few mRNA counts for these genes, we were unable to determine whether they obeyed our proposed distribution or not. Moreover, our results revealed that the CSZIP distribution exhibited a better fit with the data than the CSP distribution when focusing on a fixed time point alone, suggesting that the data are indeed zero-inflated.

**Table 1:** The proposed sample-specific hypothesis test results on whether the number of new mRNA molecules in the Cell Cycle dataset obeys the CSP and CSZIP distributions. UTD means that it is unable to determine because there are too few groupings resulting in zero degrees of freedom, when it is always a perfect fit. The significance level is 0.05.

| Labeling duration |  | 15mins | 30mins | 45mins | 60mins | 120mins | 180mins |
| --- | --- | --- | --- | --- | --- | --- | --- |
| CSP | Accept | 0.116 | 0.067 | 0.049 | 0.062 | 0.064 | 0.065 |
|  | Reject | 0.278 | 0.568 | 0.655 | 0.652 | 0.695 | 0.725 |
|  | UTD | 0.606 | 0.365 | 0.296 | 0.286 | 0.241 | 0.210 |
| CSZIP | Accept | **0.351** | **0.467** | **0.472** | **0.476** | **0.459** | **0.459** |
|  | Reject | 0.055 | 0.189 | 0.266 | 0.274 | 0.327 | 0.344 |
|  | UTD | 0.594 | 0.344 | 0.262 | 0.250 | 0.214 | 0.197 |

We next showed the high goodness-of-fit of the CSP and CSZIP model on two characteristic genes, namely *RPL41* and *IL22RA1* with an overall low and high gene expression respectively (**Fig. 3A**). Qualitatively, we found that the expected counts of both the CSP and CSZIP models matched well with the observed counts for the gene *RPL41*. Quantitatively, the results of the cell-specific chi-square test also showed that the distribution of CSP or CSZIP was well satisfies in most labeling durations (**Fig. 3A, first row**). Similar results were observed for the gene *IL22RA1* with significantly higher expression (**Fig. 3A, second row**). Therefore, we demonstrated CSP and CSZIP distribution accurately describes these two genes and is thus suitable for modeling the tscRNA-seq datasets.

**Figure 3:**
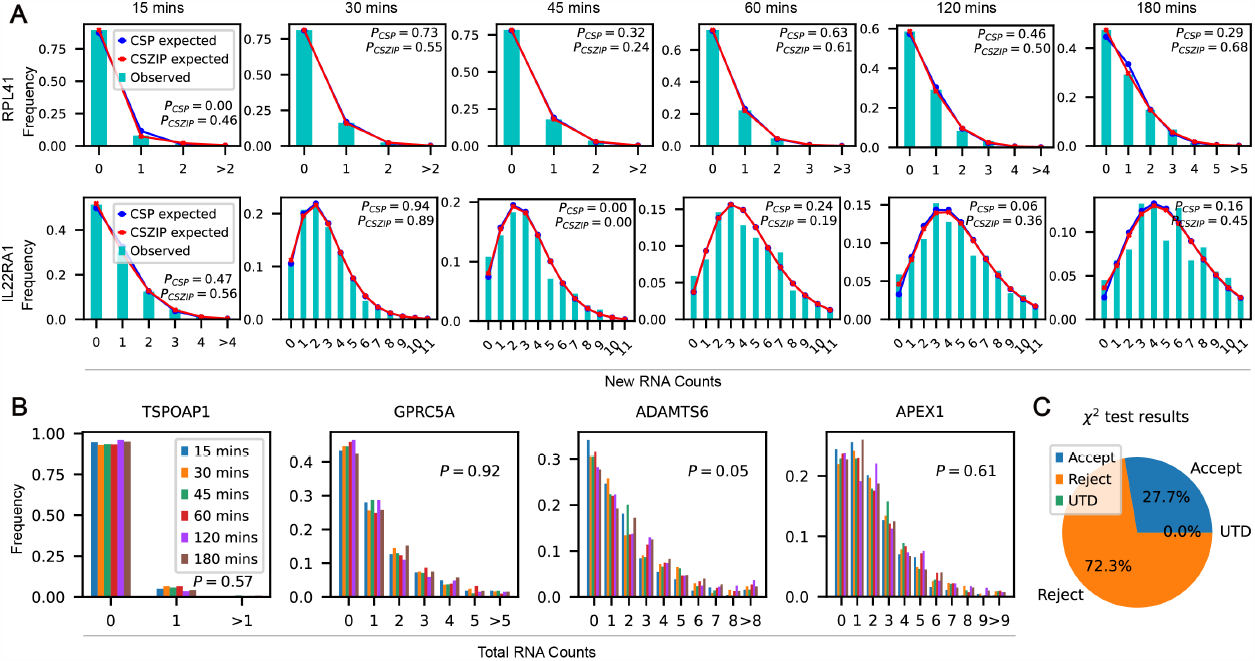
Statistical analysis of cell cycle dataset. **A**. Observed counts, expected counts of CSP, and expected counts of CSZIP of new mRNA molecules of the two example genes *RPL41* and *IL22RA1*. The first row: Fitting results of the *RPL41* gene with a small number of mRNA molecules; The second row: Fitting results of the *IL22RA1* gene with a higher number of new mRNA molecules (truncated to 11 for better visualization). **B**. Comparison of the total mRNA counts with different labeling durations of the four example genes *TSPOAP1, GPRC5A, ADAMTS6* and *APEX1*. **C**. Results of chi-square independence test for total RNA counts (significance level 0.05).

Finally, we found that, for most genes, the number of total mRNA molecules shares the same distribution across different labeling durations. In **Fig. 3B**, we showed the number of total mRNA molecules of four example genes *TSPOAP1, GPRC5A, ADAMTS6* and *APEX1* is nearly identical across different labeling durations. Quantitatively, we performed a global chi-square independence test on the number of total mRNAs (as distinct from the new mRNAs) with different durations of labeling in all genes and found that, interestingly, there are 72.3% of the genes passed the test at a significance level of 0.05 (**Fig. 3C**). This indicates that a considerable proportion of the number of genes’ total mRNA molecules obeyed the same distribution, consistent with what we observed for the four example genes.

### Storm accurately infers kinetic parameters that leads to rich insights of cell cycle via enrichment analysis

In the kinetic experiments, we relied on three stochastic models without the steady-state assumption to infer different set of kinetic parameters using maximum likelihood estimation (see “Methods section), namely *α* and *γ*_*t*_ for Model 1, *α, β* and *γ*_*s*_ for Model 2, and *α, γ*_*t*_ and *p*_off_ for Model 3. In addition, we defined the goodness-of-fit of each of the three models by utilizing the concept of deviance *R*^2^ commonly used in generalized linear models Menard (2000) (see “Methods” section). According to the goodness-of-fit index, we selected genes that were more consistent with the model assumptions for downstream tasks, such as the enrichment analysis and RNA velocity analysis, etc.

Compared with Dynamo Qiu et al. (2022), the state-of-the-art method for processing tscRNA-seq datasets, our advantages are mainly in the following aspects: (1) Our method does not require steady-state assumptions on the kinetics experiments while Dynamo heavily relies on the steady-state assumptions; (2) Our stochastic model-based approach is more consistent with real biological process, while Dynamo only utilizes the deterministic model of mean value; (3) Our model takes into account all cells in the inference, while the approach based on steady-state assumptions in Dynamo only considers a small number of cells with high expression. In addition, we revealed the difference between the total mRNA degradation rate *γ*_*t*_ and spliced mRNA degradation rate *γ*_*s*_ based on their different physical roles, distinguished them in different models, and finally gave the relationship between these two (see “Methods” section). We noted that in Dynamo, to infer *β, γ*_*t*_ was first inferred when the splicing was ignored, then 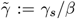 was inferred using the method based on the steady-state assumption in scVelo Bergen et al. (2020), and finally 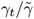 was taken as the inference of *β* upon assuming *γ*_*t*_ = *γ*_*s*_. However, 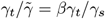, while *γ*_*t*_ and *γ*_*s*_ are generally not equal. This point was overlooked in Dynamo, which causes a inaccurate estimate of *β*. In fact, under the steady-state assumption, *β* can be directly estimated by using only *u*_*l*_(*t*) through the formula *u*_*l*_(*t*) = (1 *e*^−*βt*^)*α/β*, similar to the two-step method used in Dynamo to estimate *γ*_*t*_ through 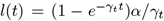 since they have similar form. However, we don’t use this method in Storm.

With the above inference methods and insights, we studied a cell cycle dataset from the scEU-seq study Battich et al. (2020). We compared the parameter inference results of the three models (**Fig. 4A**). When splicing was not considered, the inference results based on CSP and CSZIP distributions were close, with high correlation coefficients, especially in genes with higher goodness of fit (**Fig. 4A Left**). However, whether or not splicing is considered significantly impacts the inference results. The inference results based on CSP and ICSP distribution were quite different, with low correlation coefficients, even in genes with higher goodness of fit (**Fig. 4A Middle**). We speculate that this is due to the assumptions of the two models are incompatible: in CSP, *γ*_*t*_ is assumed to be a constant; while in ICSP, *γ*_*s*_ is assumed to be a constant. But these two assumptions can not be held simultaneously for their different roles in the physical modeling and our analysis results (see “Methods” section). We also compared *γ*_*t*_ and *γ*_*s*_ computed by the ICSP model, and the results showed that *γ*_*s*_ was always greater than *γ*_*t*_, and the linear correlation between the two was not high (**Fig. 4A Right**). In summary, we showed that kinetic parameters inferred from CSP and CSZIP but not CSP and ICSP, are consistent.

**Figure 4:**
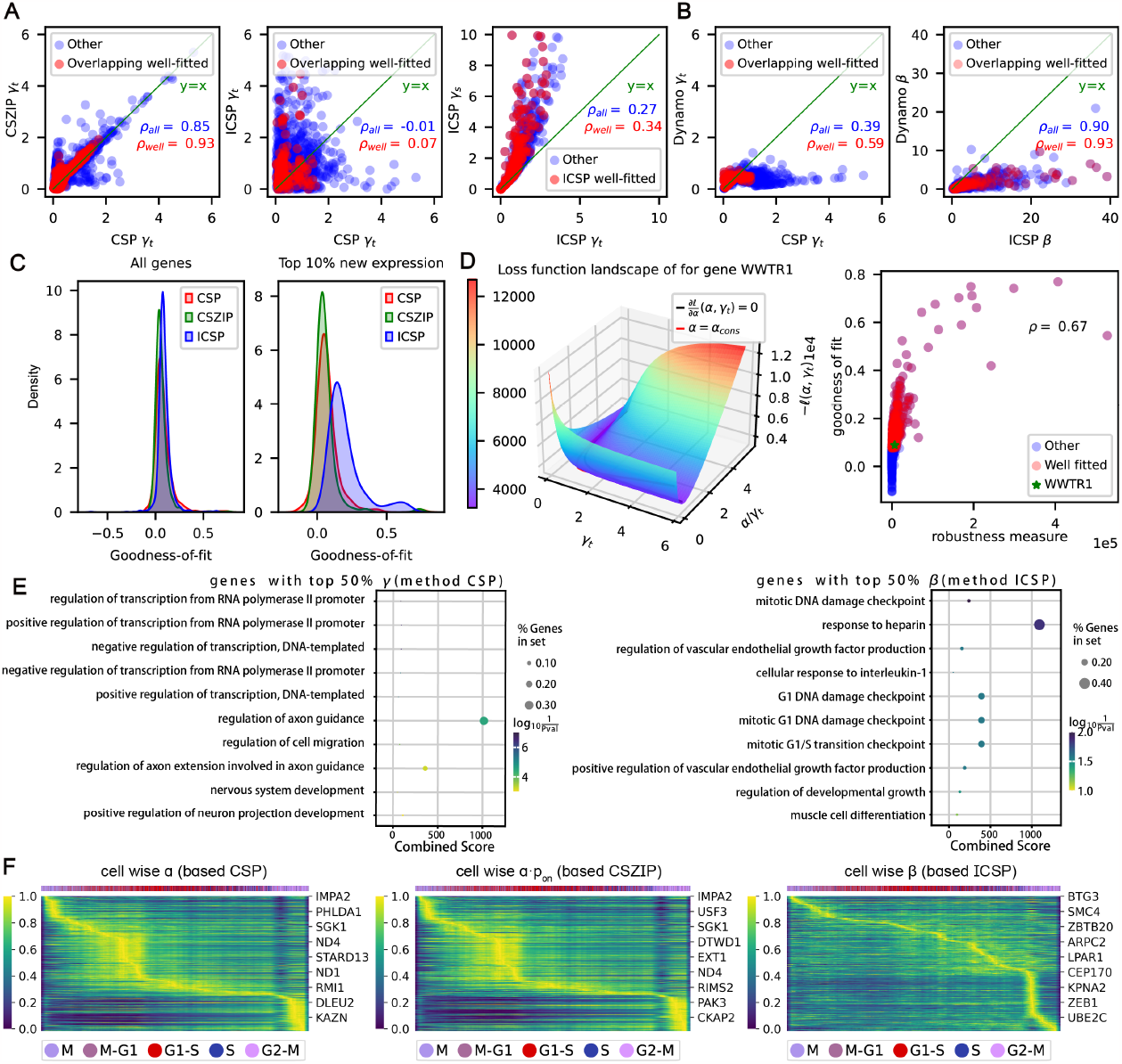
Parameter inference and enrichment analysis for the cell cycle dataset. **A**. Comparison of parameter inference results of our three stochastic models. From left to right are the comparison of *γ*_*t*_ of CSP and CSZIP, the comparison of *γ*_*t*_ of CSP and ICSP, the comparison of *γ*_*t*_ and *γ*_*s*_ in ICSP. The overlapping well-fitted genes were set as the overlap set of genes in the top 40% of the goodness-of-fit for both methods. **B**. Comparison of inferred parameters between our stochastic models and Dynamo’s method. **Left**: the comparison of *γ*_*t*_ between CSP and Dynamo. **Right**: the comparison of *β* between ICSP and Dynamo. **C**. Comparison of the goodness-of-fit of the three stochastic models. **Left**: all highly variable genes. **Right**: genes in the top 10% of average new mRNA expression in highly variable genes. **D**. Robust analysis. **Left**: Landscape of Model 1-based loss functions for the a typical gene *WWTR1*. **Right**: Scatter plot of robustness measure and goodness of fit for parameter inference. **E**. Enrichment analysis results of genes with high *γ*_*t*_, *β* (top 50%) in well fitted genes (top 40% of goodness of fit). **F**. Heat map of cell-wise parameters for well-fitted genes. From left to right, cell-wise *α* based on the CSP model, cell-wise *αp*_on_ based on the CSZIP model and cell-wise *β* based on the ICSP model, respectively. Across all three heatmaps, the X-axis is the relative cell cycle position while the order of genes in the y-axis is arranged such that the peak time of each gene increases from the top left to bottom right.

The inferred total mRNA degradation rates *γ*_*t*_ from Storm and Dynamo are close in well-fitted genes, while ICSP’s inferred splicing rates *β* are always larger than from Dynamo. We compared the inferred results of *γ*_*t*_ in our CSP model with those in Dynamo (**Fig. 4B Left**). Although they were not consistent for some genes, they are quite consistent for the genes with better fitting. We also compared the inference of *β* in our ICSP model with those in Dynamo (**Fig. 4B Right**). The result shows that the inferred *β* by our approach was usually larger than those in Dynamo, even for the genes with a better fitting. A possible explanation is that the inference of Dynamo ignored the difference between *γ*_*t*_ and *γ*_*s*_, which made the inferred *β* smaller. We also compared the goodness-of-fit of the three stochastic models. Overall, they are relatively close (**Fig. 4C Left**). However, when we focused on genes with higher new mRNA levels (top 10%), the ICSP model had a better fit (**Fig. 4C Right**). We speculate that this is because genes with higher expression are suitable to be fitted with more complex models.

When the parameter *γ*_*t*_ is small, parameter inference may not be robust enough. However, we found that the genes selected by the goodness-of-fit have robust results. We analyzed the robustness of the parameter inference in the simplest CSP model. When *γ*_*t*_*t* is small, 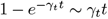 holds, then

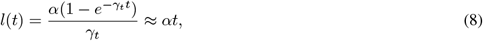

which implies that from the mean perspective the nonlinear fitting of *α* and *γ*_*t*_ degenerated into a linear fitting of *α* at this point. For a more precise analysis, let 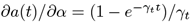, we have *∂𝓁*(*α, γ*_*t*_)*/∂α* = 0 is equivalent to

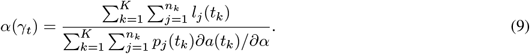

But when 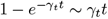 holds, *∂a*(*t*)*/∂α* ≈ *t*, then we have

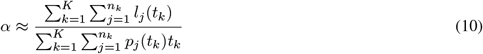

is a constant, which we denoted by *α*_cons_. We plotted the landscape of a typical negative log-likelihood loss function based on CSP model for gene *WWTR1* (**Fig. 4D Left**), with the black line corresponding to *∂𝓁*(*α, γ*_*t*_)*/∂α* = 0 (i.e. Eq. (9)) and blue line corresponding to *α* = *α*_cons_ (i.e., Eq. (10)). The landscape of the loss function shows a fairly flat area around *∂𝓁/∂α* = 0, and the two lines almost coincide when *γ*_*t*_ is small, which is consistent with our previous argument. In addition, to quantitatively measure the robustness of inference on *γ*_*t*_, since the optimal parameter is always located where the gradient is zero, we defined the *l*_1_-norm of the derivative of the loss function with respect to *γ*_*t*_ restricted to *∂𝓁/∂α* = 0 (i.e. black line),

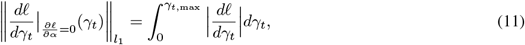

as a measure of robustness. Since the half-life of the total mRNA molecules is usually not less than half an hour, we took *γ*_*t*,max_ = 1.5. We analyzed the relationship between the robustness measure and the goodness-of-fit 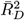 (**Fig. 4D Right**). We found that parameter robustness was positively correlated with the goodness of fit and the correlation coefficient was as high as 0.69. Though the reason for this high correlation is not clearly understood in theory, we can utilize this fact to select the genes with high goodness of fit for downstream analysis, which also ensures the results are relatively robust.

We selected the well-fitted genes (top 40% 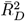) and performed enrichment analysis on this fraction according to the magnitude of gene-wise parameters *γ*_*t*_, *β, α* and *p*_off_ (**Fig. 4E, Fig. S2**). The results of the enrichment analysis showed that these genes were highly correlated with the cell cycle progression.

The assumption of constant coefficients is often violated because of the time-dependent kinetics and multiple lineages Bergen et al. (2021). Many works relaxed the constant coefficient assumption and inferred cell-specific parameters to overcome this issue Cui et al. (2022); Qiu et al. (2022); Gayoso et al. (2022); Li et al. (2023). In our proposal, we take a post-processing step to get the cell-specific parameters after inferring all parameters through previous procedures. We relaxed the constant coefficient assumption and proposed a method to infer cell-specific parameters except the constant degradation rate *γ*_*t*_ or *γ*_*s*_, i.e., we inferred cell-specific *α* in Model 1, cell-specific *α p*_on_ in Model 3, and the cell-specific *α* and *β* in Model 2 (see “Methods“ section). This partial constant coefficient assumption had support from the study in Battich et al. (2020), which showed that the degradation rate of most genes was independent of time. Finally, We plotted heat maps of the cell-wise *α* (based on CSP model), *α × p*_on_ (based on CSZIP model) and *β* (based on ICSP model) for the well-fitted genes (**Fig. 4F**). The results show that cells in the same cell cycle phase usually have closer kinetic parameters.

### Storm improves the robustness and accuracy of time-resolved RNA velocity analysis

Our three stochastic models described the evolution of the PMF (or joint PMF) of the number of new mRNA (or new unspliced and spliced mRNA) molecules over time for different settings. To estimate RNA velocity of single cells, only the evolution of the mean value over time will be considered, which requires us to reduce the stochastic models to the corresponding deterministic models (see “Methods“ section).

Based on the deterministic model derived for the mean corresponding to the three stochastic models, we inferred the relevant parameters for computing different types of RNA velocity for different models. In Models 1 and 3, we computed the total RNA velocity 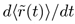 because the splicing process was ignored. In Model 2, we calculated both total RNA velocity 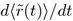 and spliced RNA velocity 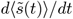 (see “Methods“ section). Note that because the new RNA velocity mostly reflects the metabolic labeling process of RNA and does not reveal RNA biogenesis, it is thus not used. In addition, a derived relationship between *γ*_*t*_ and *γ*_*s*_ suggests that the total RNA velocity can be computed based on either 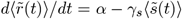 or 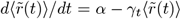. In practice, we used the former approach by default.

We compared the streamlines of the total RNA velocity of our three models with that of Dynamo on the cell cycle scEU-seq dataset (**Fig. 5A**). Almost all streamlines from our models correctly reflect the cell cycle progression, except that part of them from the ICSP model had a minor flaw in the M phase and CSZIP in the S phase. In addition, we found both ICSP and Dynamo’s spliced RNA velocity (**Fig. 5B**) did not get entirely correct streamline results. The streamlines of our ICSP model were problematic in the M-G1 phase, while the streamlines of Dynamo were problematic in the S phase. We speculate that this is probably due to the fact that new unspliced mRNAs have rather low expression levels, frustrated with many dropouts and very sparse data, resulting in unreliable inferences of the parameter *β* and inaccurate RNA velocities.

**Figure 5:**
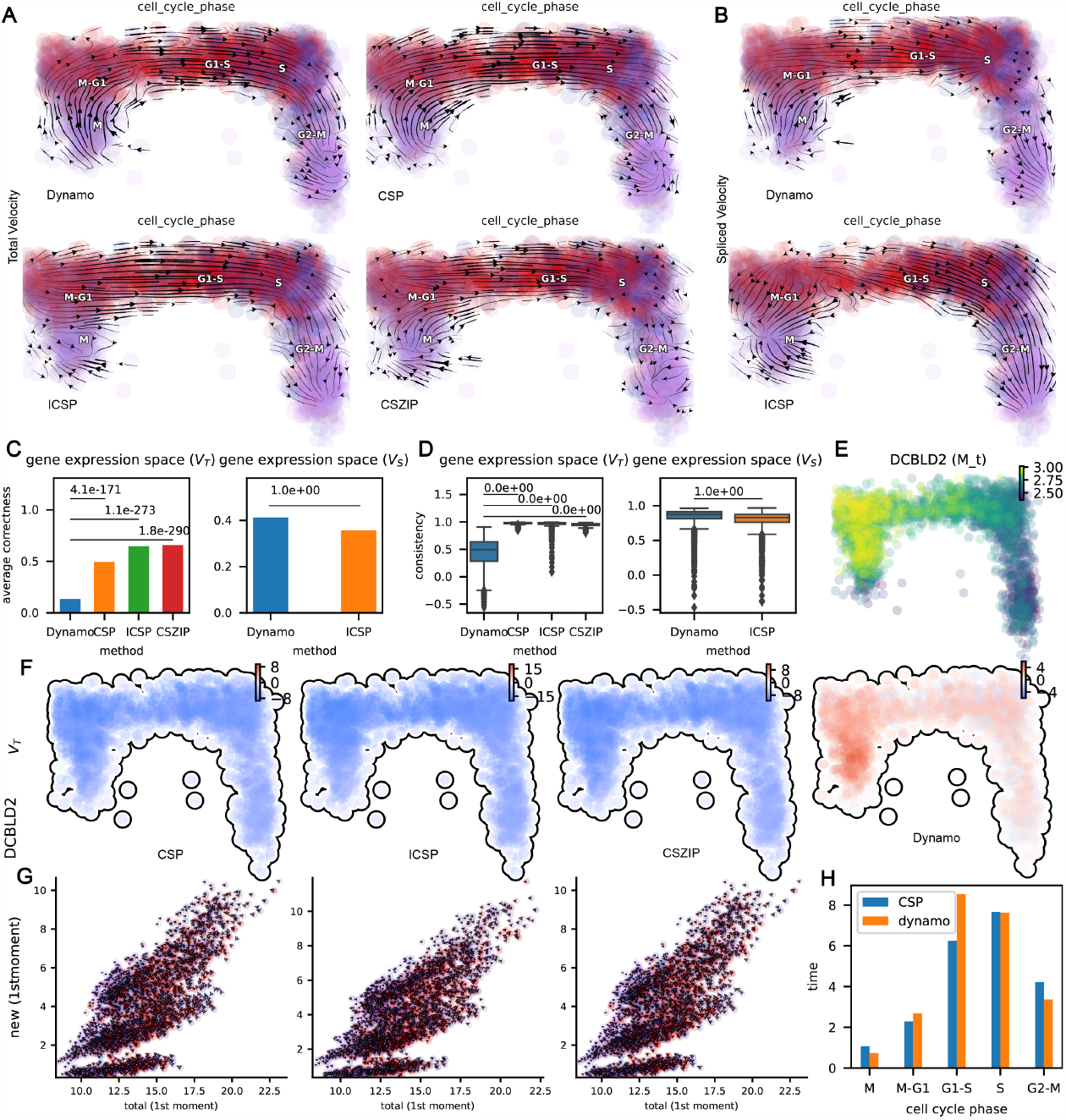
RNA velocity analysis of the cell cycle dataset. **A**. Comparison of total RNA velocity streamline visualizations between three stochastic methods and Dynamo. **B**. Comparison of spliced RNA velocity streamline visualizations between ICSP and Dynamo. **C**. Comparison of average correctness of velocity in gene expression space. **Left**: total RNA velocity. **Right**: spliced RNA velocity. The p-values are given by the one-sided Wilcoxon test. **D**. Similar to **C**, comparison of velocity consistency. **E**. The smoothed expression of *DCBLD2* in different cells. **F**. Comparison of total RNA velocity in *DCBLD2* between three stochastic models and Dynamo. **G**. Phase portraits of new-total RNA planes of *DCBLD2*. Quivers correspond to the total (x-component) or new (y-component) RNA velocity calculated by the different methods. **H**. The duration time (unit: hour) of each cell cycle phase of the human RPE1-FUCCI system based on Storm’s CSP model and Dynamo.

We also quantitatively benchmarked the average correctness and consistency of the velocities in different methods in the original gene expression space and low-dimensional space (here the RFP_GFP space is used which corresponds to the Geminin-GFP and Cdt1-RFP-corrected signals of RPE1-FUCCI cells)(**Fig. 5C,D**; **Fig. S3A,B**). The definition of correctness and consistency of velocity is given in the “Methods“ section. In the gene expression space, the average correctness and consistency of the total RNA velocity of CSP, ICSP, and CSZIP are significantly better than that of Dynamo (**Fig. 5C, D Left**), while the spliced RNA velocity of ICSP has slightly lower consistency than that of Dynamo (**Fig. 5C, D Right**). In the RFP_GFP space, the average correctness of total RNA velocity of all methods are significantly higher compared to that in the gene expression space, and simpler methods tend to improve more. The average correctness of CSP is highest at this time (**Fig. S3A Left**). However, the average correctness of the ICSP’s spliced RNA velocity still perform slightly worse than Dynamo’s (**Fig. S3A Right**). In contrast, the total RNA velocity consistency of CSP and ICSP is significantly better than that of Dynamo (**Fig. S3B Left**) and the spliced RNA velocity consistency of ICSP is also significantly better than that of Dynamo (**Fig. S3B Right**). Overall, the CSP-based total RNA velocity had the highest average correctness and consistency, significantly outperforms Dynamo, while the ICSP-based spliced RNA velocity was close to Dynamo quantitatively.

We now illustrate the advantages of our method in the estimation of kinetic parameters and the calculation of RNA velocity with two example genes: *DCBLD2* and *HIPK2*. In gene *DCBLD2*, the cells at M and M-G1 have the highest overall expression and the correct RNA velocity should be negative (**Fig 5E**). However, Dynamo returned the positive velocity, which is problematic (**Fig. 5F last column**). In contrast, CSP, CSZIP and ICSP all returned negative velocities (the **first three columns** in **Fig. 5F**). We speculated one possible explanation is that the expression of the gene *DCBLD2* has not yet reached a steady state. Consistent results were also observed from phase portraits of new-total RNA planes of *DCBLD2* (**Fig. 5G, Fig. S3C**). For gene *HIPK2*, similarly, cells in phase M and M-G1 have the highest expression overall and the correct velocity should be negative (**Fig. S3F**), but Dynamo and CSP both returned positive velocities while CSZIP got the correct results (**Fig. S3D**,**E**). We speculated one possible explanation for this is that the expression switch plays an important role in *HIPK2*.

To demonstrate the value of using gene-cell-wise parameters (except degradation rates), we visualized the streamlines of total RNA velocity based on gene-cell-wise parameters and those based only on gene-wise parameters (**Fig. S3G**). We observed that the streamlines of the CSP model and the CSZIP model in the S to G2-M phase are incorrectly reversed (**Fig. S3G Left** and **Middle**), and the streamlines of the ICSP model are also less smooth and accurate than those when gene-cell-wise parameters are used (**Fig. S3G Right**).

Finally, to demonstrate the significance of inferring time-resolved velocities with physical units, we calculated the duration time of each cell cycle phase of the human RPE1-FUCCI system based on the total RNA velocities (see “Methods“ section, **Fig. 5H**). Indeed, the human RPE1-FUCCI system has a cell-cycle time of about 21 hours (about 6 hours for G1-S phase, 8 hours for S phase, 4 hours for G2-M phase, 1 hour for M phase and 2 hours for M-G1 phase) Chao et al. (2019).

## Discussion

Storm utilizes three stochastic models for the dynamical description of new mRNAs and allows the estimation of the RNA velocity for kinetics experiments without the need for the steady-state assumption. It can also generally handle one-shot data when the steady-state assumption is enforced. One possible limitation of our model is that it does not fully utilize the total mRNA information in kinetics experiments. According to the results of the chi-square independence test, the number of total mRNA molecules of most genes obeys the same distribution. Noting that the old mRNA molecules with a labeling duration of zero are the total mRNA molecules, we think that it is a feasible direction to establish the stochastic dynamics of old mRNA and use the Wasserstein distance in optimal transport approach Vallender (1974); Zhang et al. (2021) to measure the differences between discrete distributions. Therefore, the optimal transport modeling of old RNAs may be integrated with Storm to obtain more robust RNA velocity inference. In addition, it is also worth exploring stochastic models that consider switching of gene expression states, transcription in the active state, splicing and spliced mRNA degradation simultaneously (i.e., integration of Model 2 and Model 3).

Some recent works, such as MultiVelo Li et al. (2022), Chromatin Velocity Tedesco et al. (2022), and protaccel Gorin et al. (2020), extend RNA velocity to multi-omics. It is expected that the combination of metabolic labeling technology with other multi-omics measurements will bring new opportunities, which allows for simpler parameter inference and more accurate results.

Finally, most of the existing methods make the independent gene expression assumption, and do not consider the regulatory mechanism between genes. Deep neural network approaches are promising to solve this problem. This will be an important future direction.

## Conclusions

We present Storm for estimating absolute kinetic parameters and inferring the time-resolved RNA velocity of metabolic labeling scRNA-seq data by incorporating the transient stochastic dynamics of gene expressions. Storm establishes three stochastic models of new mRNA which take into account both biological noise and cell-specific technical noise, and makes inference to the gene-specific degradation rates and other gene-cell-specific parameters without relying on the steady-state assumption in kinetics experiments. It can also handle one-shot data when the steady-state assumption is adopted. Numerical results show that Storm is able to accurately fit the kinetic cell cycle dataset and many one-shot experimental datasets. In addition, our numerical experience suggests that Model 1 (i.e., the CSP model) outperforms the other two models when splicing dynamics is not of interest, and the Model 2 (i.e., the ICSP model) is the valid choice if the data contains both labeling and splicing information and splicing dynamics is of interest. However, further applications and performance evaluations for more challenging datasets with temporal information are desired and it will be studied in the future.

## Methods

### Derivation of three stochastic dynamical models

Here we developed three stochastic models for the dynamical description of new mRNAs: Model 1) a stochastic dynamical model of new mRNA involving only metabolic-labeling transcription and degradation; Model 2) a stochastic dynamical model of new unspliced and spliced mRNA involving metabolic-labeling transcription, splicing and spliced mRNA degradation; and Model 3) a stochastic dynamical model of new mRNA involving gene state switching, metabolic-labeling transcription and degradation.

### Model 1: Stochastic dynamical modeling of new mRNA

Following Battich et al. (2020); Qiu et al. (2022), we made the following assumptions: (1) Genes are independent. (2) Both the transcription rate *α* and the degradation rate of total mRNA *γ*_*t*_ are constants.

The chemical master equation (CME) for the new/labeled mRNA 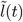, corresponding to the chemical reactions shown in **the first row of Fig. 1A**, is given by

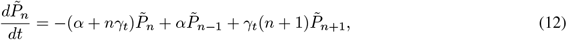

where 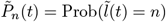. The initial value of new mRNA count is zero, i.e., 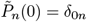, where

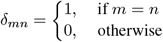

is the Kroneckers delta function. The solution of Eq. (12) is

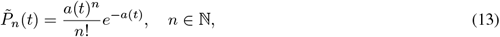

where 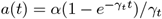. This means that 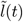 obeys the Poisson distribution with mean *a*(*t*).

The above stochastic model only describes the true expression count of new mRNA 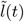 in a cell with labeling duration *t*, but the measured sequencing data is different from this count due to technical noise. Denote by *l*(*t*) the number of measured new mRNA molecules, and assume that *l*(*t*) is associated with 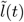 through a dropout process, which we modeled as a binomial distribution:

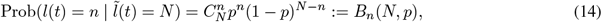

where *p* is the capture probability of a single mRNA molecule. We further assume that the total number of mRNA molecules across all genes in different cells are close, which was commonly adopted in the preprocessing step La Manno et al. (2018); Bergen et al. (2020); Qiu et al. (2022). Denote by *n*_*j*_ the total number of mRNA molecules across all genes in cell *j*, i.e., *n*_*j*_ = Σ_*i*_ *r*_*ij*_, where *r*_*ij*_ refers to the number of mRNA molecules in gene *i* of cell *j* in the scRNA-seq measurements. This assumption implies that the capture probability of mRNA molecules in different cells is different, and *p*_*j*_ ∝ *n*_*j*_. In our computation, we took *p*_*j*_ = *n*_*j*_ */n*_med_, where *n*_med_ is the median of *n*_*j*_.

We denoted the PMF of new mRNA sequencing result *l*_*j*_ (*t*) of cell *j* with labeling duration *t* by

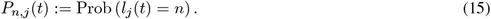

Then

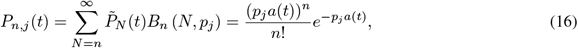

which means that *l*_*j*_ (*t*) obeys the Poisson distribution with mean *p*_*j*_ *a*(*t*).

In summary, the former derivation shows that the number of new mRNA molecules in different cells in scRNA-seq measurements obeys Poisson distribution with cell-specific parameters, and these parameters were proportional to *p*_*j*_, i.e., proportional to *n*_*j*_. We call this distribution the *cell-specific Poisson distribution*.

### Model 2: Stochastic dynamical modeling of new unspliced and spliced mRNAs

Compared with Model 1, we distinguished whether an mRNA molecule is spliced or not and incorporated the splicing process, which was shown in **the first row of Fig. 1A**. Again we assumed that the genes are independent. In addition, we further assumed that the transcription rate *α*, splicing rate *β*, and spliced mRNA degradation rate *γ*_*s*_ are all constants.

The CME for the new/labeled unspliced and spliced mRNAs 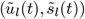, corresponding to the considered chemical reactions shown in **the first row of Fig. 1A**, is given by

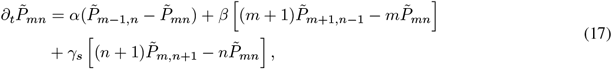

where 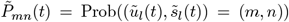. The initial distribution of new unspliced and spliced mRNA is 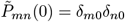. The solution of Eq. (17) is

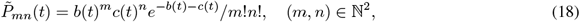

where

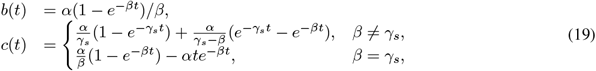

which means that 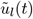 and 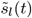 obey independent Poisson distributions with mean *b*(*t*) and *c*(*t*), respectively. We refer interested readers to Li et al. (2021) for derivation details.

Denote by (*u*_*l*_(*t*), *s*_*l*_(*t*)) the number of measured new unspliced and spliced mRNA molecules in the scRNA-seq experiments with labeling duration *t*. By assuming that the dropout processes for new unspliced and spliced mRNAs are independent and the capture probability is independent of whether they are spliced or not, we modeled the dropout process for 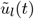 and 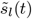 as independent binomial distributions with the same parameter *p*. So we got

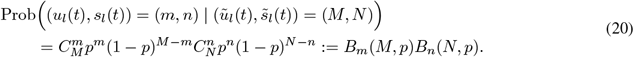

For the same reason as Model 1, we take *p*_*j*_ proportional to *n*_*j*_. And we took *p*_*j*_ = *n*_*j*_ */n*_med_ in the computation.

We denoted the joint PMF of new unspliced and spliced mRNA sequencing counts (*u*_*l,j*_ (*t*), *s*_*l,j*_ (*t*)) of cell *j* with labeling duration *t* by

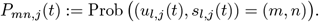

Then

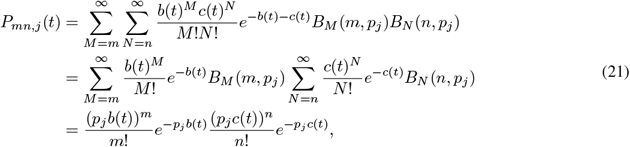

which means that *u*_*l,j*_ (*t*) and *s*_*l,j*_ (*t*) are independently Poisson distributed with mean *p*_*j*_ *b*(*t*) and *p*_*j*_ *c*(*t*), respectively.

In summary, (*u*_*l*_(*t*), *s*_*l*_(*t*)) obeys independent cell-specific Poisson distribution.

### Model 3: Stochastic dynamical modeling of new mRNA considering switching

In Model 3, we further considered the on/off gene state switching shown in **the first row of Fig. 1C**. We assumed that the genes are independent as well, and the transcription rate *α*, mRNA degradation rate *γ*_*t*_, the gene on-to-off rate *k*_off_ and off-to-on rate *k*_on_ are all constants. Furthermore, we assumed that *k*_on_ and *k*_off_ are significantly smaller than *α* and *γ*_*t*_, which implies that the gene expression is either always on or always off during the transcription/degradation period. From Eq. (12), it is known that cells in the on state obey a Poisson distribution with mean *a*(*t*), while cells in the off state do not express. Define *p*_off_ = *k*_off_ */*(*k*_off_ + *k*_off_). Then 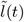 obeys the zero-inflated Poisson distribution

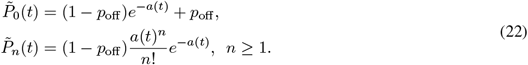

Similarly, by taking into account the technical noise in scRNA-seq experiments, the PMF of *l*_*j*_ (*t*) is

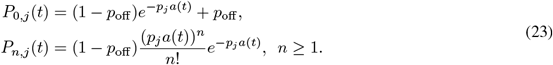

In summary, different cells obey the ZIP distribution with different parameters as shown in Eq. (23), which we called cell-specific zero-inflated Poisson distribution.

### Chi-square goodness-of-fit test for cell-specific distributions at a fixed time

We would construct an asymptotic *χ*^2^ statistic for the data with common distribution type but sample-specific parameters. This goodness-of-fit test is to assess whether the null hypothesis that the considered data, at a fixed labeling duration, obeys the proposed distribution can be accepted.

We first divided the value range of the considered data into *c* classes. According to the range that the samples fall in, we got *n* independent categorically distributed random samples *X*_*i*_ ∈ {1, 2, …, *c*} for *i* = 1, 2, …, *n* with sample dependent parameter *p*_*i*_, respectively. An equivalent representation for the categorical variable *X*_*i*_ is to denote *X*_*i*_ = (*X*_*ij*_)_*j*=1,…,*c*_ ∈{*e*_1_, …, *e*_*c*_}, where *e*_*j*_ = (*δ*_*jk*_)_*k*=1,…,*c*_ is the indicator vector for *j* = 1, …, *c*. Correspondingly, the parameter *p*_*i*_ = (*p*_*i*1_, …, *p*_*ic*_)^*T*^ is a *c*-dimensional vector with non-negative elements and sums to one, which is defined as

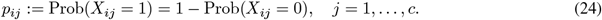

This implies that Var(*X*_*ij*_) = *p*_*ij*_ (1 − *p*_*ij*_) and Cov(*X*_*ij*_, *X*_*il*_) = 𝔼 [*X*_*ij*_ *X*_*il*_] −*p*_*j*_ *p*_*l*_ = −*p*_*j*_ *p*_*l*_ for *j* ≠ *l*. Therefore, the covariance matrix of random vector *X*_*i*_ is

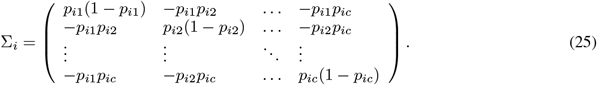

For sample *i*, we defined the truncated random vector 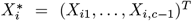 and truncated vector 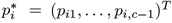, which is the first *c* − 1 components of *X*_*i*_ and *p*_*i*_, respectively. The covariance matrix of 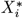 is the submatrix consisting of the upper-left (*c* − 1) × (*c* − 1) block of Σ_*i*_, denoted by 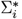, which can be written as

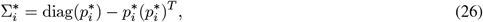

where 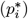 is the diagonal matrix formed by the components of 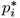.

Define 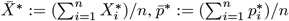 and 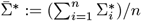 and let

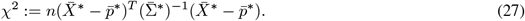

Below we would show that *χ*^2^ is an asymptotic chi-square statistic with degrees of freedom *c* − 1. First note that

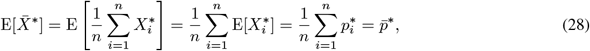

then the covariance

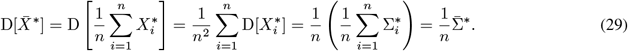

Let 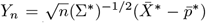. When *n* goes to infinity, *Y*_*n*_ converges in distribution to the normal distribution *N* (0, *I*_*c*−1_) according to the central limit theorem for the independent sum of random variables. Thus, 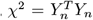 converges in distribution to a chi-square distribution with degrees of freedom *c* − 1.

In summary, we proposed a new asymptotic *χ*^2^ statistic for sample-specific distributions. For a fixed labeling duration *t*_fixed_, *a*(*t*_fixed_), *b*(*t*_fixed_) and *c*(*t*_fixed_) are all constants, the proposed *χ*^2^ statistics can be used to test whether the new mRNA sequencing data are consistent with the CSP, ICSP and CSZIP distributions based on Models 1, 2 and 3, respectively. In addition, since there are one, two and two parameters to be inferred in CSP, ICSP and CSZIP, respectively, the same number of degrees of freedom should be subtracted. Following Koehler and Larntz (1980), we ensured that the expected count *np*_*j*_ ≥ 0.25 in each group when determining the group value ranges. Finally, we take *p*-value as 0.05 in the computation.

### Parameter inference in one-shot experiments

In the one-shot experiments, we only observed new RNA *l*_*j*_ (*t*) and total RNA *r*_*j*_ (*t*) data for one labeling duration *t*. So we had to invoke the steady-state assumption for the total RNA in this case.

When the dynamics of total RNA in Model 1 is at steady state, i.e.,

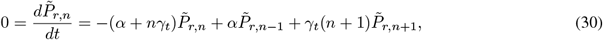

where 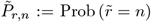 is the invariant PMF of the true expression of total RNA. From Eq. (16) we know that when technical noise is considered, the observed total RNA counts obey a similar CSP distribution

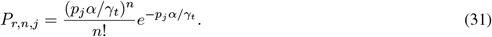

At this point, we obtained the distributions of the new RNA and total RNA observations so that parameter inference can be performed using the MLE. We want to maximize the log-likelihood function

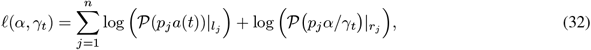

where 𝒫 (*λ*) _*n*_ := Prob(*X* = *n*) = *e*^−*λ*^*λ*^*n*^*/n*! is the probability of *X* = *n* for a Poisson-distributed random variable *X* with mean *λ*. When *∂ℓ/∂α* = 0 and *∂ℓ/∂γ*_*t*_ = 0, the likelihood function is maximized and it can be solved analytically

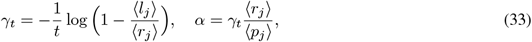

where ⟨·⟩ means the population average defined by

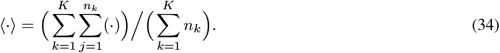

Since here it is for the one-shot data set, K = 1. Note that Eq. (33) is similar to the formula in Dynamo Qiu et al. (2022) for estimating the parameters for one-shot data. The difference is that this formula averages the raw counts, while the method in Dynamo averages the smoothed data.

### Parameter inference in kinetics experiments

In the kinetics experiments, we observed data *l*_*j*_ (*t*_*k*_) (or (*u*_*l,j*_ (*t*_*k*_), *s*_*l,j*_ (*t*_*k*_))) for new mRNA (or new unspliced and spliced mRNAs) with different labeling durations. We assumed that there are *K* labeling durations *t*_*k*_ for *k* = 1, 2, …, *K*, and the number of cells with labeling duration *t*_*k*_ is *n*_*k*_. We utilized the MLE to infer the unknown parameters in different models without relying on steady-state assumptions.

In Model 1, we need to maximize the log-likelihood function

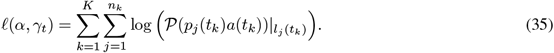

It is equivalent to minimizing the following loss function

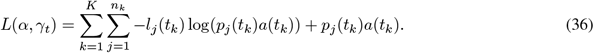

The optimum of the loss is achieved when the gradient equals 0. Utilizing the concrete expression of *a*(*t*) (Eq. (3)) in Model 1, we got 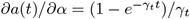. Then *∂L*(*α, γ*_*t*_)*/∂α* = 0 has a closed form solution

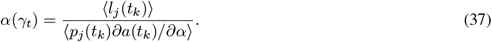

Another component of the Euler-Lagrange equation *∂L/∂γ*_*t*_ = 0 has no closed form solution, so we need to solve *γ*_*t*_ by numerical iterations. We took the initial value of *γ*_*t*_ as the solution from Dynamo Qiu et al. (2022) under the steady-state assumption. Denote it as *γ*_*t*,0_, and correspondingly, we take the initial value of *α* as *α*_0_ = *α*(*γ*_*t*,0_).

In Model 2, we need to maximize the log-likelihood function

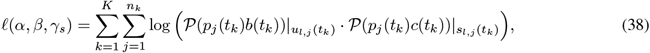

which is equivalent to minimizing the loss function

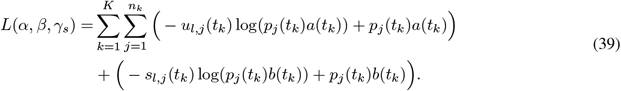

Utilizing (19), we got 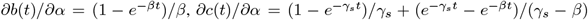 when *β* ≠ *γ*_*s*_, and the case for *β* = *γ*_*s*_ is similar. So *∂L*(*α, β*_*t*_, *γ*_*s*_)*/∂α* = 0 has a closed form solution

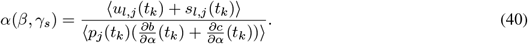

However *∂L/∂β* = 0 and *∂L/∂γ*_*s*_ = 0 have no closed form solution, and we need to solve these equations by iterations. The choice of initial values is similar to the Model 1 case. We took the initial value of *β* and *γ*_*s*_ as the solution from Dynamo Qiu et al. (2022) under the steady-state assumption, which we denoted as *β*_0_, *γ*_*s*,0_. And then the initial value of *α* is taken as *α*_0_ = *α*(*β*_0_, *γ*_*s*,0_).

In Model 3, we need to maximize the log-likelihood function

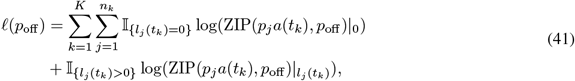

where ZIP(*λ, p*_off_) |_*n*_ := Prob(*X* = *n*) is the probability of *X* = *n* for a ZIP-distributed random variable *X* with parameters *λ* and *p*_off_. It is equivalent to minimizing the loss function

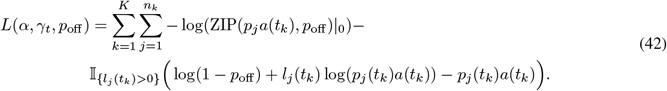

Similar as before, we chose the initial value of *γ*_*t*_, denoted as *γ*_*t*,0_, based on the steady state assumption, and chose the moment estimator

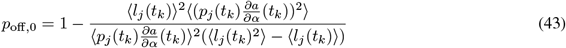

and

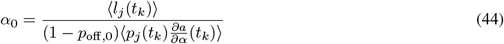

as the initial values of *p*_off_ and *α*.

According to the biological meaning of the parameters, we added the constraints 0 < *α* < 10*α*_0_, 0 < *β* < 10*β*_0_, 0 < *γ*_*t*_ < 10*γ*_*t*,0_, 0 < *γ*_*s*_ < 10*γ*_*s*,0_ and 0 < *p*_off_ < 1, and we called the SLSQP optimizer in SciPy to solve the above optimization problem.

### Goodness-of-fit test for the distribution evolution in time

In ordinary least squares (OLS) linear regression, people often use

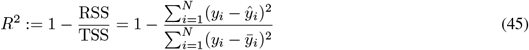

to define the goodness of fit, where *y*_*i*_ is the sample observation, *ŷ*_*i*_ is the model prediction, and 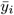 is the sample mean. For the generalized linear model (GLM), the *R*^2^ can be defined using the deviance *D* and null deviance *D*_0_ Menard (2000),

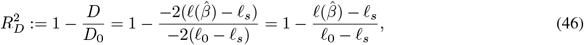

where 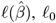 and *ℓ*_*s*_ denotes the log-likelihood function of the model with parameter 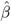, the null model (that is, fitted with only the intercept), and the saturated model (that is, fitted with one parameter per sample), respectively. A pictorial representation of *D* and *D*_0_ is shown in Fig. 1E. 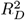 can be seen as a generalization of *R*^2^, which is equal to *R*^2^ when the model is a least squares linear regression Menard (2000). Finally, to overcome the disadvantage of adding more parameters without reducing 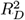 (similar to *R*^2^), we used adjusted 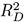 (denoted as 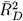) as the goodness of fit of our model, which is defined as

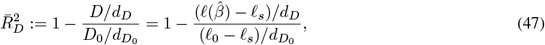

where *d*_*D*_ and 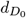 are the degrees of freedom of *D* and *D*_0_, respectively.

In Model 1, *ℓ*_*s*_ has the closed form

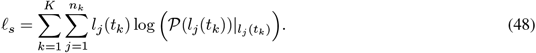

To calculate *ℓ*_0_, we need to maximize the log-likelihood function

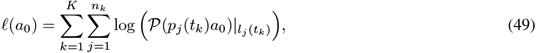

where *a*_0_ is the intercept. The problem has a closed form solution *a*_0_ = ⟨*l*_*j*_ (*t*_*k*_)⟩*/*⟨*p*_*j*_ (*t*_*k*_)⟩. In addition, *d*_*D*_ = *N* − 2 and 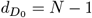, where *N* is the number of cells.

In Model 2, *ℓ*_*s*_ has the closed form

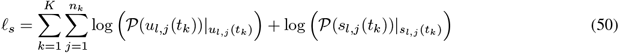

To calculate *ℓ*_0_, we need to maximize the log-likelihood function

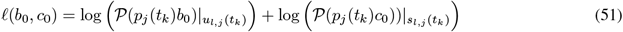

where *b*_0_ and *c*_0_ are intercepts and have closed form solutions *b*_0_ = ⟨*u*_*l,j*_ (*t*_*k*_)⟩*/*⟨*p*_*j*_ (*t*_*k*_)⟩ and *c*_0_ = ⟨*s*_*l,j*_ (*t*_*k*_)⟩*/*⟨*p*_*j*_ (*t*_*k*_)⟩, respectively. In addition, *d*_*D*_ = 2*N* − 3 and 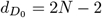

In Model 3, to calculate *ℓ*_*s*_, we need to maximize the log-likelihood function

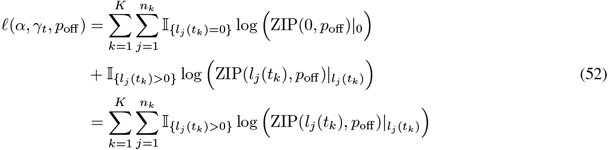

When *p*_off_ is equal to zero, Eq. (52) is maximized, and the closed form solution of *ℓ*_*s*_ is

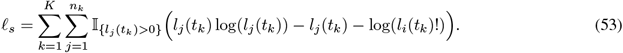

To calculate *ℓ*_0_, we need to maximize the log-likelihood function

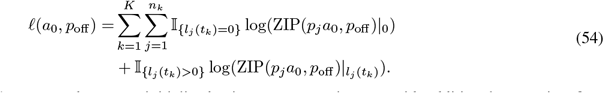

Similar to solving Eq. (42), *p*_off,0_ and *a*_0_ were initialized using moment estimators with additional constraints 0 < *p*_off_ < 1 and 0 < *a* < 10*a*_0_. We then called the SLSQP optimizer in SciPy to solve the problem. In addition, *d*_*D*_ = *N* − 2 and 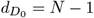.

### Post-processing for cell-specific parameters

In our cell-specific modeling of gene expression, we only assumed that *γ*_*t*_ (in Models 1 and 3) and *γ*_*s*_ (in Model 2) are constants over cells and are inferred based on the corresponding stochastic models, while the other parameters are cell-specific and continuously dependent on gene expressions. This relaxed assumption implies that only the degradation rate is common to all cells, and only cells with similar gene expressions have similar other parameters (due to continuous dependence). To realize this assumption, we first constructed the k-nearest neighbor (kNN) graph of cells by a data preprocessing. The cell-specific parameter inference was performed by applying the inference to the kNN graph for each cell with local constant parameter assumption and already inferred degradation rates. The inference details for our three models were shown as below.

In Model 1, we have

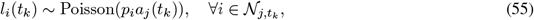

where 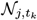 denotes the set of top *k* cells that have the most similar gene expressions as the *j*th cell with labeling duration *t*_*k*_ (including itself) and 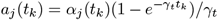. Assuming that *γ*_*t*_ has been inferred, we can obtain a local estimator

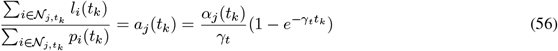

by using the MLE. Define 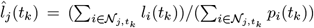, Then the cell-specific transcription rate *α*_*j*_ (*t*_*k*_) has a closed form solution

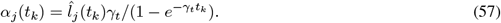

In Model 2, we have

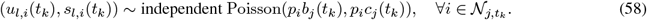

Similarly, assuming *γ*_*s*_ has been inferred, and defining the local estimators

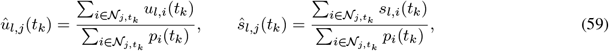

we have

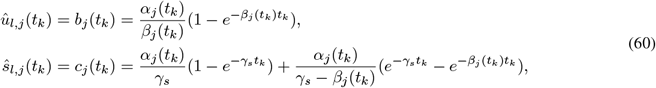

which is a nonlinear system. We have

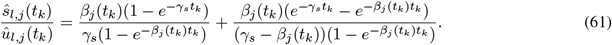

To solve *β*_*j*_ (*t*_*k*_), we set its initial value as previously inferred *β* by global constant assumption. We then call the *foot* function in SciPy to solve the nonlinear equation (61) to get *β*_*j*_ (*t*_*k*_). The *α*_*j*_ (*t*_*k*_) has a closed form solution

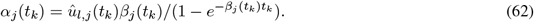

In summary, in Model 2, we can infer the cell-specific transcription rate *α*_*j*_ (*t*_*k*_) and splicing rate *β*_*j*_ (*t*_*k*_).

In Model 3, we have

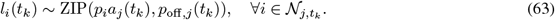

When computing RNA velocity, we only need to know *α*_*j*_ (*t*_*k*_)(1 − *p*_off,*j*_ (*t*_*k*_)) as a whole, and not their respective values (see next subsection). To simplify the computation, we used the moment estimation instead of MLE, and got

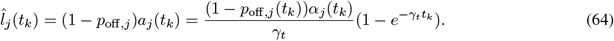

Similarly, assuming *γ*_*t*_ has been inferred, *α*_*j*_ (*t*_*k*_)(1 − *p*_off,*j*_ (*t*_*k*_)) has a closed form solution

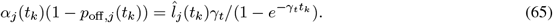

### Reduction from stochastic to deterministic models for RNA velocity

We used discrete counts data in the proposed parameter inference and goodness-of-fit calculation via stochastic models. However, when we need to compute and visualize the RNA velocity, we should take the reduction from stochastic to deterministic models to get the mean velocity. Below we would show the reduction process and reveal the connection between the stochastic and their corresponding deterministic models.

⟨ ⟩ ⟨ ⟩

In Model 1, let us denote the mean value of 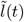 by 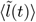, which is defined as 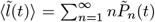. From Eq. (12) we can obtain the deterministic equation after suitable algebraic manipulations

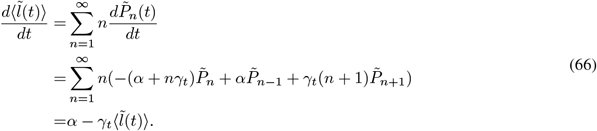

Similarly, the mean value of total RNA 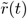 satisfies the equation

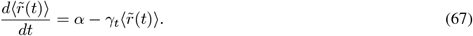

Since the initial value of 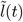 is zero, we got

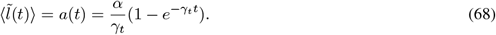

In Model 2, the marginal PMFs of 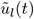 and 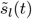 are

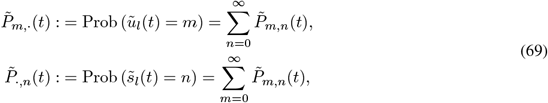

respectively. The mean values of 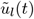 and 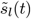 have the form 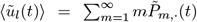 and 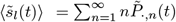. From the CME (17), we can obtain

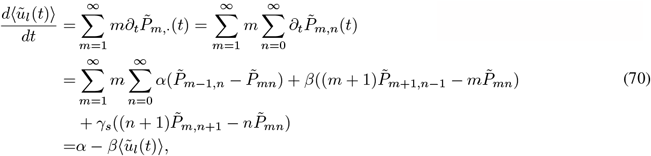

and

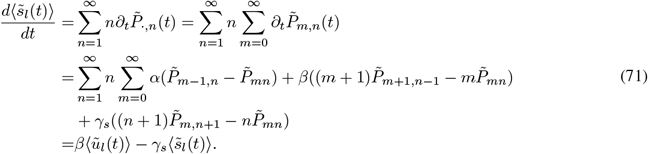

Similarly, we can derive the equations for the mean values of total unspliced and spliced mRNA 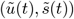:

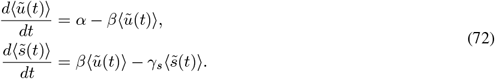

Since the initial value of 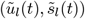 is (0, 0), we got

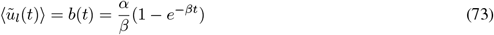

and

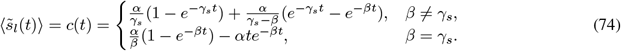

Similar to Model 1, in Model 3, 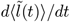 and 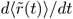 satisfy the equations

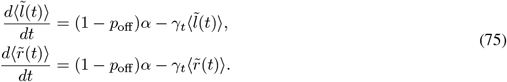

Since the initial value of 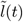 is zero, we got

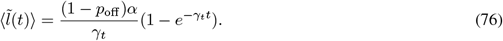

### Computation of RNA velocity

To ease the notation, we denoted the new mRNA after data preprocessing by 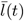, defined as

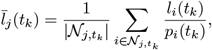

which is different from the true expression 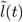, the discrete counts data *l*(*t*), and the notation 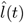 in the post-processing subsection. We would also use the notation 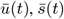 and 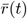 with similar definition.

In Model 1, only the total RNA velocity can be obtained due to the lack of the splicing stage. From Eq. (67), we have

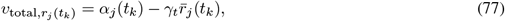

where 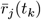 is the number of total mRNA molecules of the *j*th cell labeled with length *t*_*k*_ after data preprocessing.

In Model 2, we add the two equations in Eq. (72) to obtain

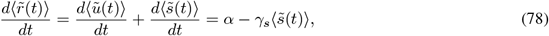

and thus get the equation for total RNA velocity

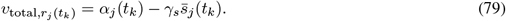

In addition, in Model 2, we can also calculate the spliced RNA velocity by the following equation

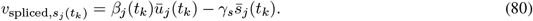

Similar to Model 1, the total RNA velocity in Model 3 can be obtained by the equation

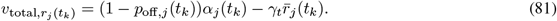

### Relationship between *γ*_*t*_ and *γ*_*s*_ and its implications

The difference between Eqs. (67) and (78) implies the difference between the total mRNA degradation rate *γ*_*t*_ and spliced mRNA degradation rate *γ*_*s*_. After suitable manipulations, we had the relation between *γ*_*t*_ and *γ*_*s*_ as below

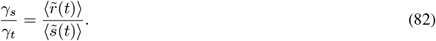

Therefore, we naturally got a method to infer *γ*_*t*_ when *γ*_*s*_ is known. Specifically, we first performed a zero-intercept linear regression

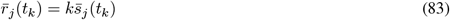

to get the slope *k*. Then we computed *γ*_*t*_ by *γ*_*t*_ = *γ*_*s*_*/k*. Therefore, we can also infer *γ*_*t*_ and compute the total RNA velocity by Eq. (77) in Model 2.

We would also like to point out that Model 1 and 3 are incompatible upon assuming that *γ*_*t*_ and *γ*_*s*_ are both constants. These two assumptions usually do not hold simultaneously. Otherwise, from Eq. (82) we knew that 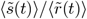 is a constant, which is equivalent to that 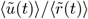 is a constant, i.e., 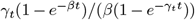 is a constant. But this is only true when *β* and *γ*_*t*_ are equal.

### Definition of correctness and consistency of velocity

The correctness of cell velocities is defined as follows: Consider the cell *i* with position *x*_*i*_ and velocity *v*_*i*_. Define its one-step extrapolated position as *x*_*i*_ + *v*_*i*_. We say that *v*_*i*_ is correct (correctness index = 1) if the cell *j* closest to the extrapolated position *x*_*i*_ + *v*_*i*_ ranks after *i* in the temporal ordering. Otherwise the correctness does not hold and we set the correctness index to be 0. Thus the average correctness refers to the percentage of correct velocities.

The consistency means the extent to which the velocity of one cell is consistent with the velocities of its neighboring cells, and we use the average cosine similarity proposed in scVelo Bergen et al. (2020) to measure this consistency.

### Calculation of cell cycle time

After the total RNA velocities are obtained, we can evaluate the time of each phase of a cell cycle based on them. Specifically, we first pick *k* cells 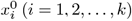 whose relative positions are closest to 0 as a cell group, calculate their average expression 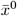 and velocity 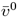 as the initial expression *x*^0^ and velocity *v*^0^, and extrapolate the state of the cell group with a short time step *dt*, that is, *x*^1^ = *x*^0^ + *v*^0^*dt*. We then search for another *k* cells 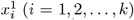 which are closest to the extrapolated state *x*^1^, set the majority of the phase of these *k* cells to the phase of *x*^1^, and set their average velocity 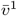 as *v*^1^ for the second cell group. Next, the extrapolation and local *k*-cells group identification step can be repeated until a given threshold of the relative position is exceeded. In the actual calculation, we set *k* = 300, *dt* = 0.01, and the threshold of the relative position to be 88% quantile of all relative positions. The above approach for processing the cell groups instead of cells themselves is to reduce the data noise by local averaging.

## Data availability

In this study, we used the following public tscRNA-seq datasets from scSLAM-seq Erhard et al. (2019), scNT-seq Qiu et al. (2020), sci-fate Cao et al. (2020) and scEU-seq Battich et al. (2020). These datasets can be downloaded directly through the Python package Dynamo.

## Code availability

Storm is implemented in Python and is available at https://github.com/aristoteleo/CSP4ML.

## Author’s contributions

TL and XQ designed the research. QP performed the research. All of the authors analyzed the data and wrote the paper.

## Acknowledgements

We thank Prof. Fang Yao for helpful discussions. TL and QP acknowledge the support from NSFC and MSTC under Grant No.s 11825102, 12288101 and 2021YFA1003300.

## Additional Files

**Additional file 1 — Supplementary information with supplemental Figures S1-S3**.

