## Supplementary information for "Storm: Incorporating transient stochastic dynamics to infer the RNA velocity with metabolic labeling information"

In this supplementary material, we present three supplemental Figures S1 to S3 which are related to Figures 2, 4 and 5 in the main text, respectively.

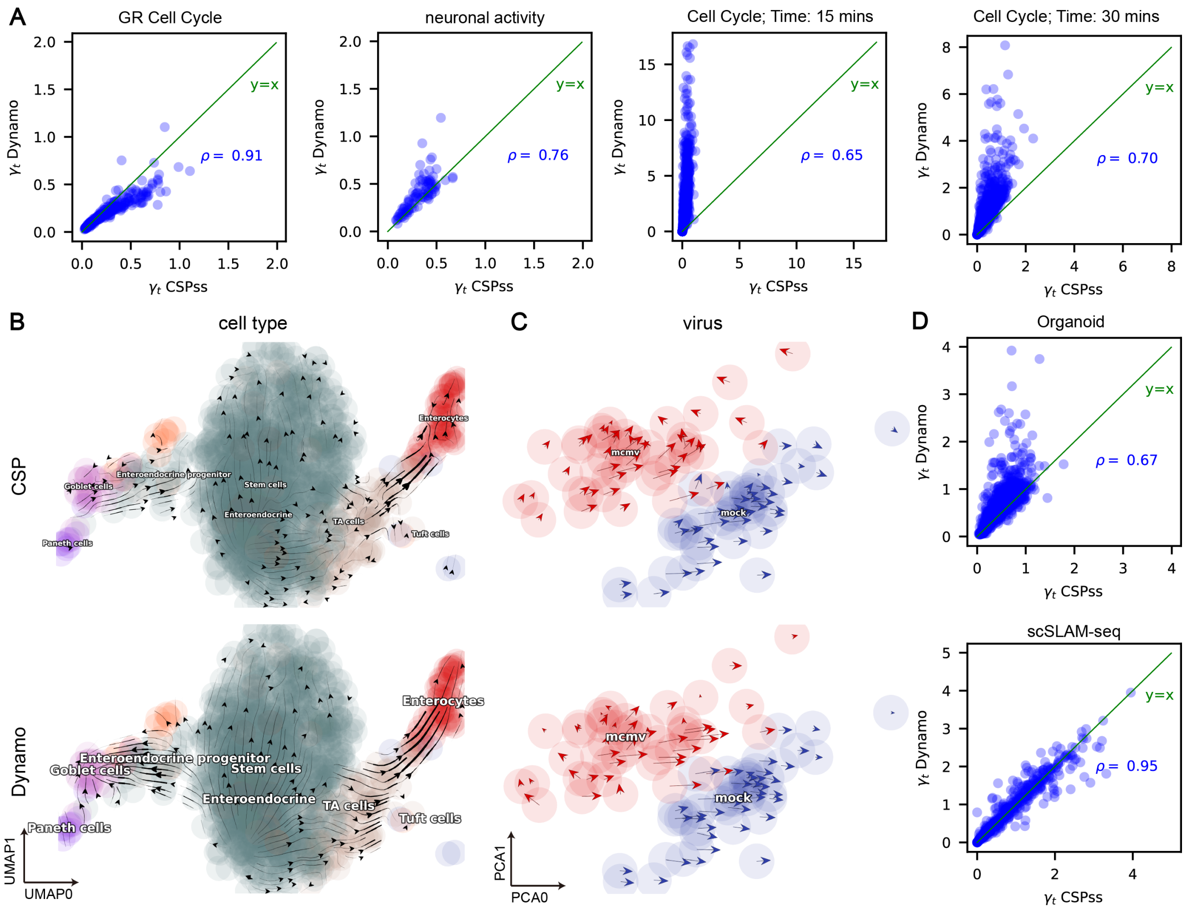


Figure S1: **Storm analyses on additional one-shot datasets, realated to Figure 2. A.** Degradation rates $\gamma_t$ estimated with our method compared to that of the Dynamo method in the sci-fate dataset [52020Cao et al.], the neuronal activity dataset from scNT-seq [242020Qiu et al.], and the cell cycle dataset from scEU-seq [12020Battich et al.] with labeling durations of 15 minutes and 30 minutes. **B.** Streamline plot in the UMAP space of the murine intestinal organoid system dataset from scEU seq [12020Battich et al.]. **C.** Cell quiver plot in the PCA space of the scSLAM-seq dataset [92019Erhard et al.]. **D.** Same as **A**, but for the organoid system dataset from scEU-seq [12020Battich et al.] and for the scSLAM-seq dataset [92019Erhard et al.].

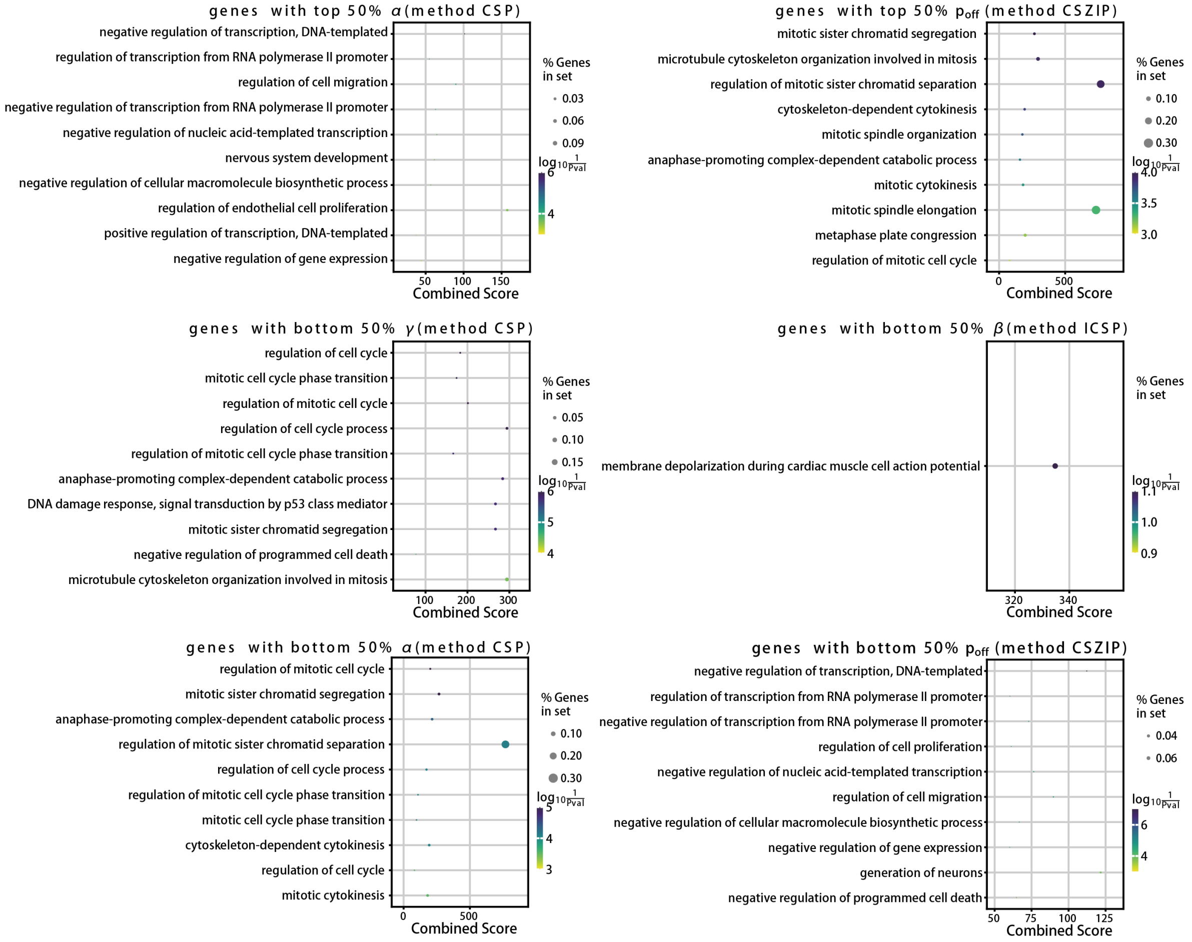


Figure S2: **GO (gene ontology) pathway enrichment results of genes with high $\alpha$ and $p_{\text{off}}$ (top 50%) and low $\gamma_t$, $\beta$, $\alpha$ and $p_{\text{off}}$ (bottom 50%) in well-fit genes (top 40% of goodness of fit), related to Figure 4 in main text.**

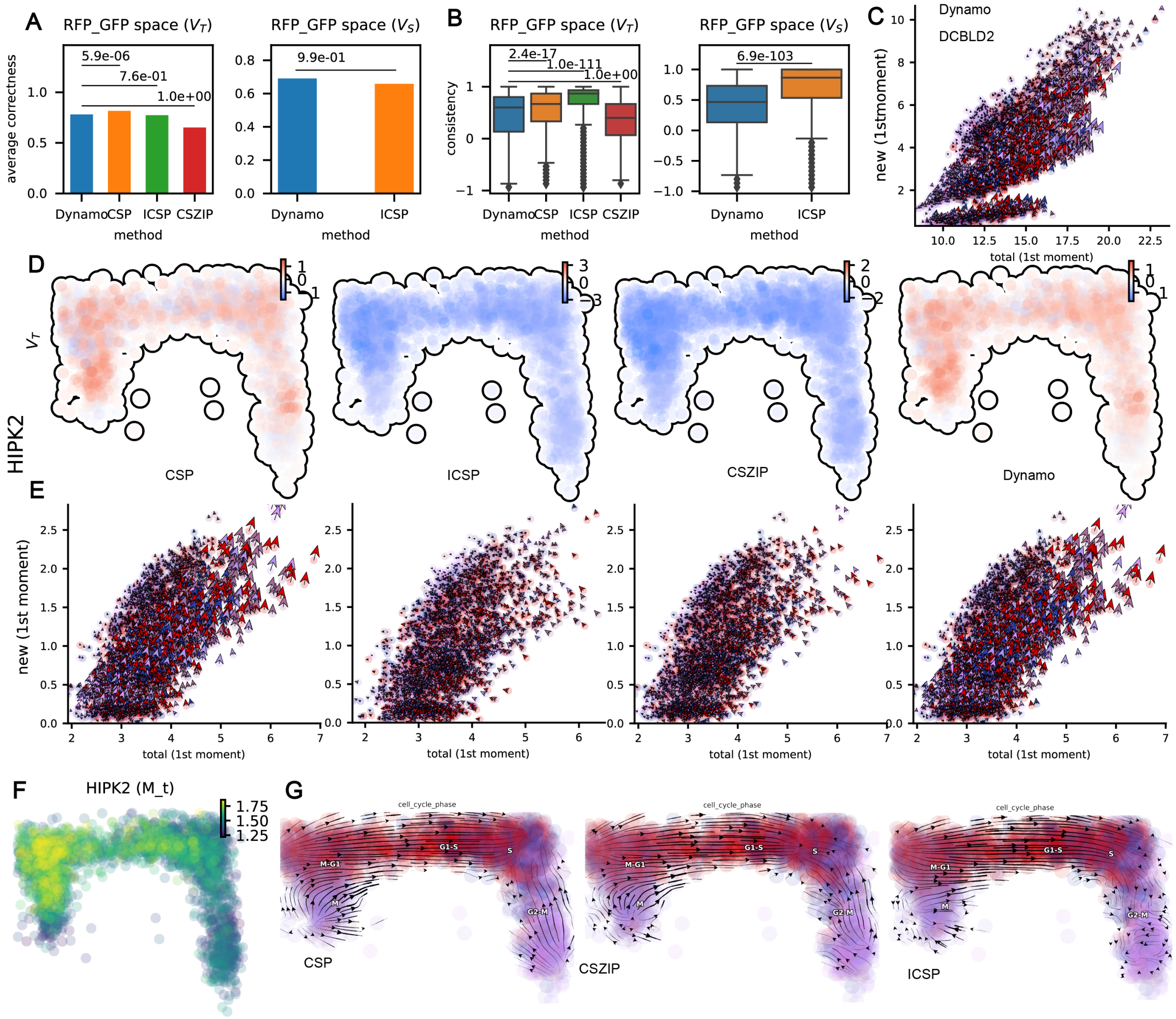


Figure S3: **RNA velocity analysis of the cell cycle dataset, related to Figure 5 A.** Comparison of the average correctness of velocity in RFP_GFP space. **Left**: total velocity. **Right**: spliced velocity. The p-values are given by the one-sided Wilcoxon test. **B.** Similar to **A**, but for velocity consistency. **C.** Phase portraits of new-total RNA planes of *DCBLD2* of Dynamo. Quivers correspond to the total (x-component) or new (y-component) RNA velocity calculated by the different methods. **D.** Comparison of total RNA velocity in *HIPK2* between three stochastic methods and Dynamo. **E.** Similar to **C**, but for gene *HIPK2* of three stochastic methods and Dynamo. **F.** The smoothed expression pattern of *HIPK2* across cells. **G.** Total RNA velocity streamlines calculated using gene-wise parameters (instead of using gene-cell-wise parameters except for the degradation rate). **Left**: CSP. **Middle**: CZISP. **Right**: ICSP
